# Spike mutations that affect the function and antigenicity of recent KP.3.1.1-like SARS-CoV-2 variants

**DOI:** 10.1101/2025.08.18.671001

**Authors:** Bernadeta Dadonaite, Sheri Harari, Brendan B. Larsen, Lucas Kampman, Alex Harteloo, Anna Elias-Warren, Helen Y. Chu, Jesse D. Bloom

**Affiliations:** Basic Sciences Division and Computational Biology Program, Fred Hutchinson Cancer Center, Seattle, Washington, 98109, USA; Molecular and Cellular Biology Graduate Program, University of Washington, Seattle, WA; University of Washington, Department of Medicine, Division of Allergy and Infectious Diseases, Seattle, WA; Howard Hughes Medical Institute, Seattle, WA, 98195, USA

## Abstract

SARS-CoV-2 is under strong evolutionary selection to acquire mutations in its spike protein that reduce neutralization by human polyclonal antibodies. Here we use pseudovirus-based deep mutational scanning to measure how mutations to the spike from the recent KP.3.1.1 SARS-CoV-2 strain affect cell entry, binding to ACE2 receptor, RBD up/down motion, and neutralization by human sera and clinically relevant antibodies. The spike mutations that most affect serum antibody neutralization sometimes differ between sera collected before versus after recent vaccination or infection, indicating these exposures shift the neutralization immunodominance hierarchy. The sites where mutations cause the greatest reduction in neutralization by post-vaccination or infection sera include receptor-binding domain (RBD) sites 475, 478 and 487, all of which have mutated in recent SARS-CoV-2 variants. Multiple mutations outside the RBD affect sera neutralization as strongly as any RBD mutations by modulating RBD up/down movement. Some sites that affect RBD up/down movement have mutated in recent SARS-CoV-2 variants. Finally, we measure how spike mutations affect neutralization by three clinically relevant SARS-CoV-2 antibodies: VYD222, BD55-1205, and SA55. Overall, these results illuminate the current constraints and pressures shaping SARS-CoV-2 evolution, and can help with efforts to forecast possible future antigenic changes that may impact vaccines or clinical antibodies.

**Importance:** This study measures how mutations to the spike of a SARS-CoV-2 variant that circulated in early 2025 affect its function and recognition by both the polyclonal antibodies produced by the human immune system and monoclonal antibodies used as prophylactics. These measurements are made with a pseudovirus system that enables safe study of viral protein mutations using virions that can only infect cells once. The study identifies mutations that decrease recognition by current human antibody immunity; many of these mutations are increasingly being observed in new viral variants. It also shows the importance of mutations that move the spike’s receptor binding domain up or down. Overall, these results are useful for forecasting viral evolution and assessing which newly emerging variants have reduced recognition by immunity and antibody prophylactics.

## Introduction

Over the course of SARS-CoV-2 evolution in humans over the last half decade, the spike protein has accumulated >60 amino-acid mutations (1–3). This evolution is driven by strong selective pressure for spike to escape from the antibody immunity accumulating in the human population (4–7) while retaining its ability to bind ACE2 receptor (8,9) and mediate cell entry (10,11). New SARS-CoV-2 lineages carrying additional mutations in spike are constantly emerging, but it remains challenging to predict which of these lineages have mutations that will enable them to be evolutionary successful.

Deep mutational scanning is a powerful approach to measure how spike mutations affect key functional and antigenic properties of spike (2,9,12–14), but the fact that both spike (8,15,16) and human population immunity (17–20) are constantly evolving limit the utility of measurements made using older strains and human antibodies for understanding newer variants. Here, we use pseudovirus-based deep mutational scanning (2,21) to measure how thousands of mutations to the spike of the recent KP.3.1.1 variant affect cell entry, receptor binding, RBD up/down motion, and neutralization by human sera and therapeutic antibodies. Overall, our work provides detailed maps of the functional and antigenic effects of spike mutations that can help rationalize recent trends in SARS-CoV-2 evolution and identify mutations that affect key protein properties.

## Results

### Pseudovirus-based deep mutational scanning of KP.3.1.1 spike

To measure how mutations in the SARS-CoV-2 spike affect cell entry, receptor binding and escape polyclonal sera or therapeutic antibodies, we used pseudovirus-based deep mutational scanning (**Fig. 1A**) (2,21). This method produces genotype-phenotype linked lentiviral particles that encode uniquely barcoded spike variants and can be used to measure the effects of mutations on different spike phenotypes (21) (**Fig. S1A**). Because these pseudoviruses are restricted to a single round of infection and require helper plasmids to produce viral particles, they cannot cause disease or transmit in humans, making them a safe tool for characterising mutations in viral proteins at biosafety-level-2.

**Figure 1.**
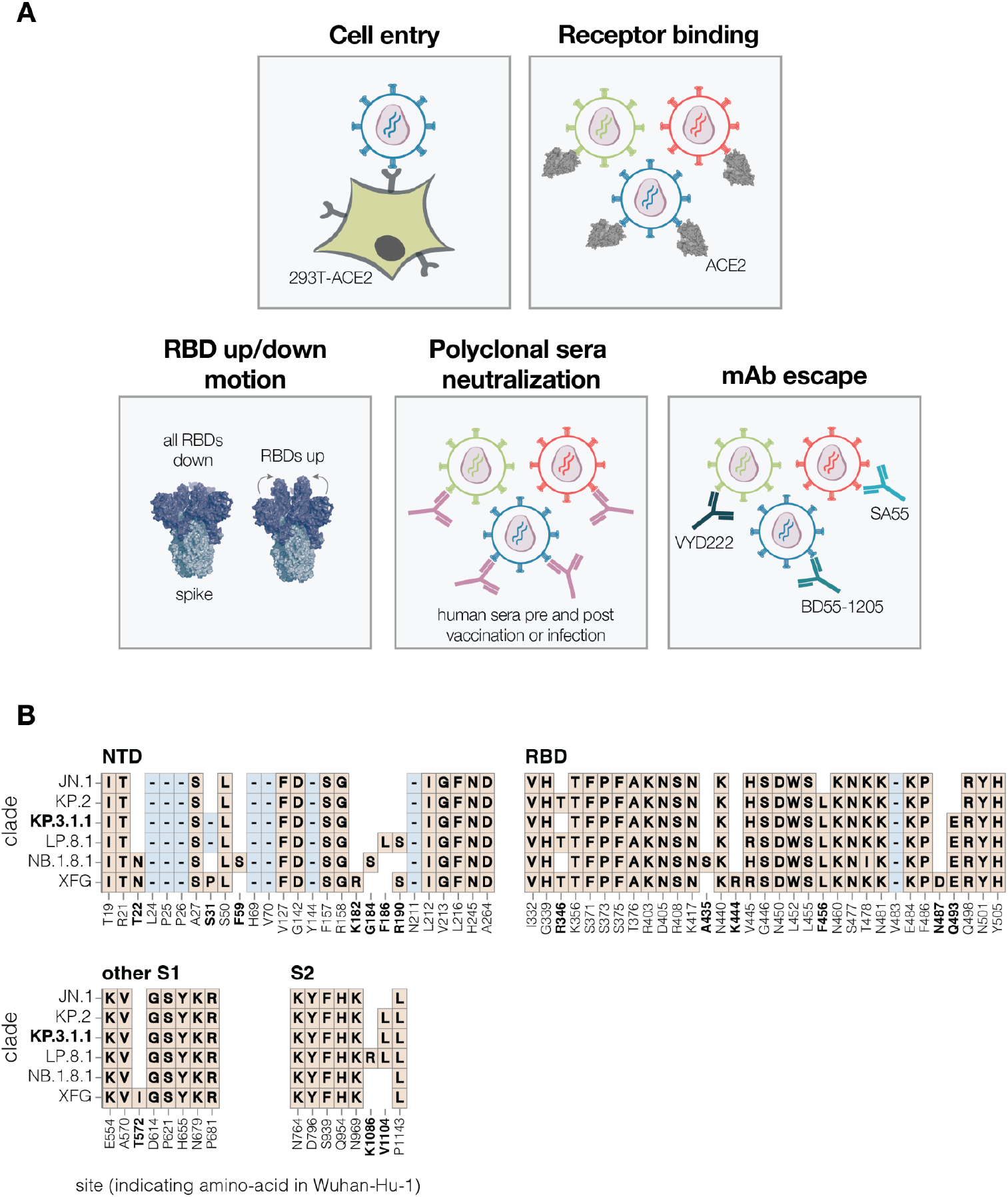
Measurement of different spike phenotypes using KP.3.1.1 spike deep mutational scanning. **A**. We measured the effects of mutations in the KP.3.1.1 spike on pseudovirus entry into 293T cells expressing ACE2, binding to ACE2 receptor, RBD up/down motion, neutralization by human sera, and neutralization by several key monoclonal antibodies. **B**. Spike amino-acid mutations and deletions in the KP.3.1.1 spike used in our deep mutational scanning and other key JN.1-descendant lineages relative to the early Wuhan-Hu-1 strain. Site labels indicate the amino-acid identity and residue number in the Wuhan-Hu-1 strain. Sites that differ among JN.1 and its descendant strains are bolded; non-bolded sites have fixed mutations relative to Wuhan-Hu-1 shared among all the lineages shown. When a variant has the same identity at a site as Wuhan-Hu-1, this is indicated with empty white space. Insertions are not shown; all JN.1 descendant lineages have an MPLF amino-acid insertion at position 16.

We designed pseudovirus-based deep mutational scanning libraries for the spike protein from the recently circulating KP.3.1.1 strain. KP.3.1.1 is a descendant of the JN.1 lineage and was one of the major variants circulating from the second half of 2024 to early 2025 (22). Its spike shares many important antigenic mutations with the other current JN.1 descendant strains, and is closely related to the spikes currently recommended as options for inclusion in SARS-CoV-2 vaccines (JN.1, KP.2, and LP.8.1) (**Fig. 1B**) (23).

We designed the deep mutational scanning libraries to contain all evolutionarily accessible and antigenically important mutations in the spike protein. Specifically, we included all mutations that have occurred at appreciable frequency during the SARS-CoV-2 evolution in humans, as well as every possible amino-acid change at sites that have mutated frequently in recent variants and all sites within the RBD. We produced two independent pseudovirus libraries (Lib-1 and Lib-2), which contained 42,783 and 45,513 barcoded variants, respectively, and covered 95% of the 9,809 targeted amino-acid mutations with an average of 1.3 mutations per spike (**Fig. S1B-C**).

### Mutation effects on spike-mediated cell entry

We measured how mutations to KP.3.1.1 spike affect entry into 293T cells that were engineered to express medium levels of the ACE2 receptor (24) (**Fig. 2A** and interactive heat map at https://dms-vep.org/SARS-CoV-2_KP.3.1.1_spike_DMS/cell_entry.html). The measured effects of mutations on cell entry were highly correlated between the two independent libraries (**Fig. S1D**). As expected stop codons were highly deleterious for cell entry whereas amino acid mutations had varied effects (**Fig. 2A**). Single-residue deletions were well tolerated at many sites in the N-terminus domain (NTD), consistent with frequent NTD deletions in many circulating SARS-CoV-2 variants (25) (**Fig. 2A**). Amino-acid mutations in the RBD had a range of effects, with some sites intolerant of mutations but others tolerant of many changes. Note that the measurements of mutation effects on cell entry here were made using only a single cell line and some mutations may have different effects on cell entry in other cell lines due to differences in for example receptor or protease expression (26,27).

**Figure 2.**
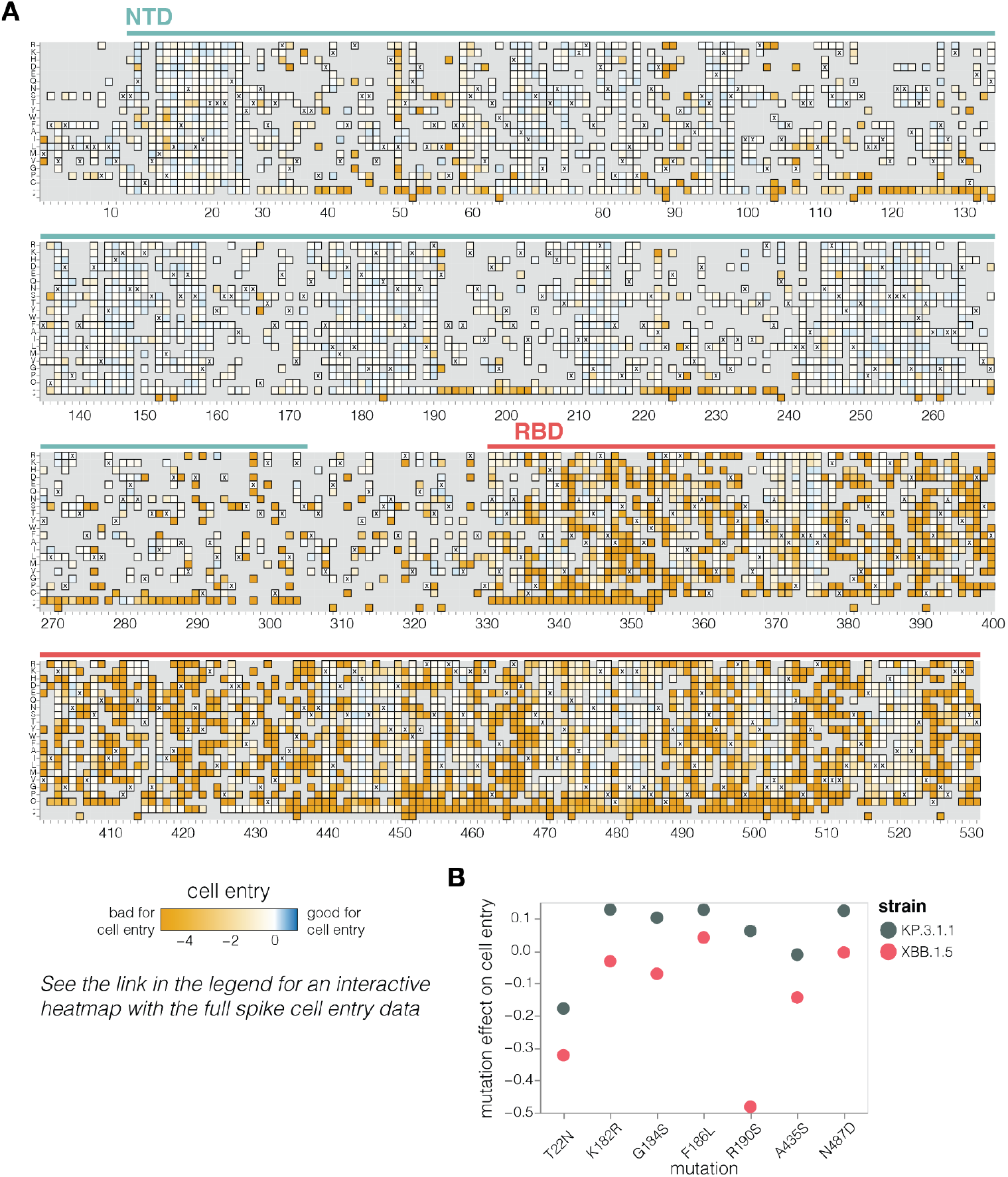
Effects of mutations to the KP.3.1.1 spike on pseudovirus entry in ACE2-expressing cells. **A**. Effects of mutations in spike on entry in 293T cells expressing a medium amount of ACE2 (24). Effects greater than zero (blue) indicate a mutation improves cell entry while effects less than zero (orange) indicate a mutation impairs cell entry. X indicates the wild-type amino acid in KP.3.1.1. Light grey indicates mutations for which effects were not measured in our libraries; note that our library design excluded most mutations expected to be highly deleterious from all regions of the spike except for the RBD. Due to space constraints this figure shows only the NTD and RBD; see https://dms-vep.org/SARS-CoV-2_KP.3.1.1_spike_DMS/cell_entry.html for an interactive heatmap that shows mutations across the full spike. **B**. Effects on cell entry for some key recent mutations in the KP.3.1.1 versus XBB.1.5 spikes. The effects in the KP.3.1.1 spike are from the current study, the effects in the XBB.1.5 spike were published previously (2).

Our measurements suggest a possible reason why certain mutations have begun to recurrently evolve in recent JN.1-descended strains related to KP.3.1.1 after being rare in earlier variants. A number of these mutations—specifically T22N, K182R, G184S, F186L, R190S, A435S, and N487D—are better tolerated for cell entry in the KP.3.1.1 spike compared to the earlier pre-JN.1 XBB.1.5 lineage (**Fig. 2B**), as assessed by comparing our current deep mutational scanning to prior measurements for the XBB.1.5 spike (2). Therefore, shifts in mutational tolerance for specific mutations may be a contributor to the recent recurrent selection for these mutations.

### Mutation effects on ACE2 binding

To determine how spike mutations affect receptor binding, we measured how well each spike mutant pseudovirus was neutralized by soluble monomeric ACE2 protein (**Fig. S2A**). We and others have previously shown that ACE2 binding affinity to spike is proportional to neutralization of SARS-CoV-2 pseudovirus by soluble ACE2 protein (2,28,29). Namely, mutations that increase spike’s binding to ACE2 also increase pseudovirus neutralization by soluble ACE2 protein, and mutations that decrease spike’s ACE2 binding decrease pseudovirus neutralization by soluble ACE2. Therefore, incubating deep mutational scanning libraries with increasing amounts of monomeric ACE2 protein allows us to measure how mutations affect ACE2 binding. Note that this approach only works for spike mutants that retain at least some moderate ability to mediate pseudovirus entry in ACE2-expressing cells. Among the spike mutations that retain sufficient cell entry function, effects on cell entry and ACE2 binding show no correlation (**Fig. S2B**), demonstrating that cell entry and ACE2 binding are distinct phenotypes, and ACE2 binding is often not the limiting factor for cell entry in our assays.

A variety of mutations both in the RBD and other regions of spike affect ACE2 binding, as measured by soluble ACE2 neutralization (**Fig. 3A-B** and interactive heatmap at https://dms-vep.org/SARS-CoV-2_KP.3.1.1_spike_DMS/receptor_binding.html). The substantial effect of some mutations outside the RBD on ACE2 binding is because the interaction of the full spike with ACE2 is impacted by several distinct mechanisms: direct interaction of the RBD with ACE2, changes in RBD up (open) or down (closed) conformation, and changes to S1 shedding (30–33). Interestingly, we measure mutations at sites distant to the RBD’s ACE2 binding motif to have as large effects on ACE2 binding as mutations at sites in close proximity to ACE2, emphasizing the importance of conformational changes to spike in affecting ACE2 binding (**Fig. 3B**). Many ACE2 distal RBD mutations with the strongest binding effects are at sites near the base of the RBD in spike, suggesting their likely involvement in positioning the RBD in the up or down conformation (eg, sites 332, 358, 390, 393, 395, 517 and 527; **Fig. 3A-B** and **Fig S2C**). Among the sites in proximity to ACE2, certain mutations at site E493 cause the largest increase in receptor binding (**Fig. 3A-B**). Notably site 493 interacts with ACE2 directly, recently substituted from Q to E in parents of KP.3.1.1 and several other current lineages, and has been previously shown to epistatically interact with two other recent mutations also present in KP.3.1.1 (L455S and F456L) (7,15).

**Figure 3.**
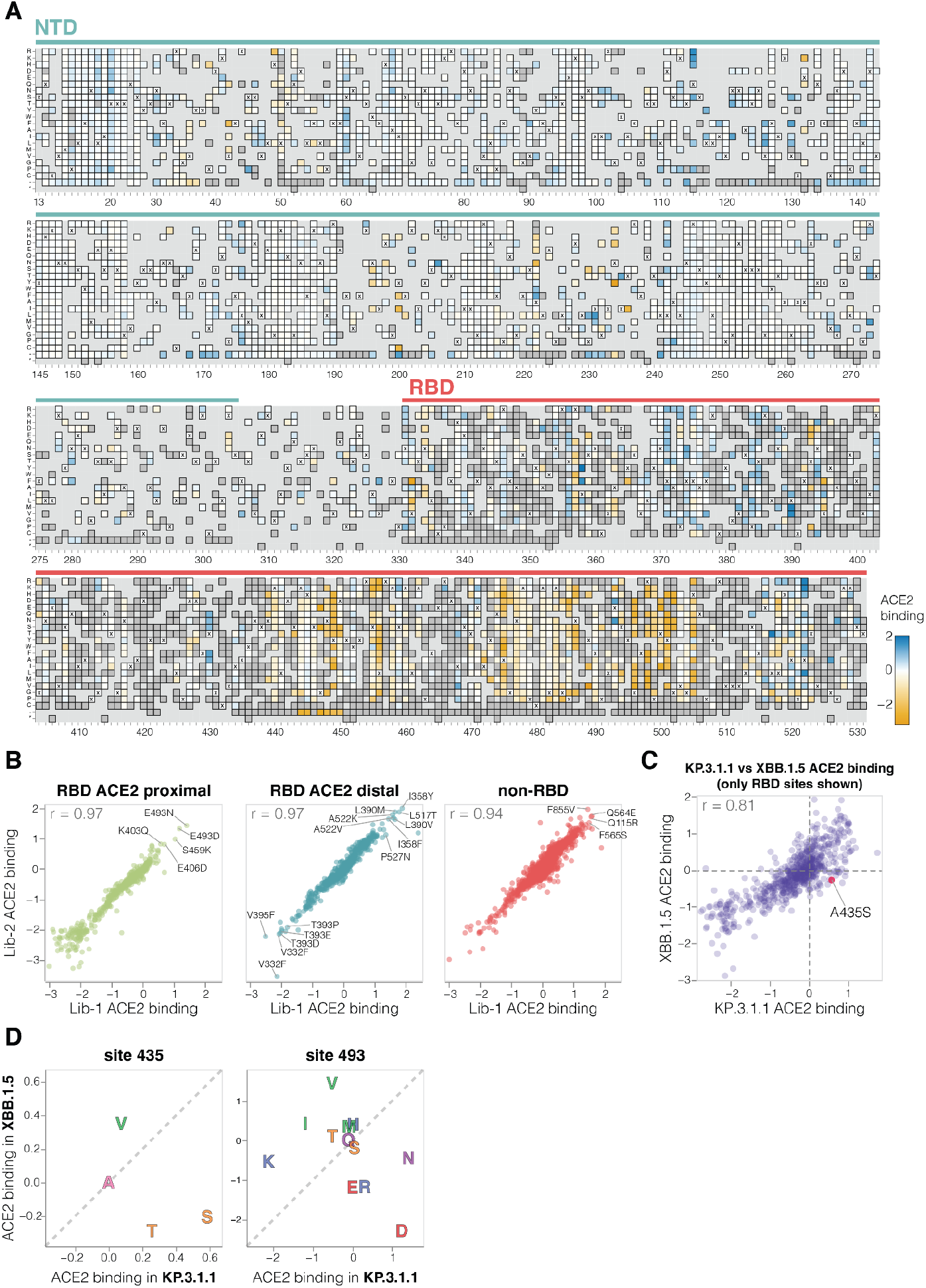
Effects of mutations to the KP.3.1.1 spike on ACE2 binding. **A**. Effects of mutations in spike on ACE2 binding. Effects greater than zero (blue) indicate a mutation improves ACE2 binding while effects less than zero (orange) indicate a mutation decreases ACE2 binding. X indicates wild-type amino acid in KP.3.1.1. Dark grey indicates mutations that were present in our libraries but too deleterious for cell entry to measure an effect on ACE2 binding; light grey indicates mutations for which effects were not measured in our libraries. Due to space constraints this figure shows only the NTD and RBD; see https://dms-vep.org/SARS-CoV-2_KP.3.1.1_spike_DMS/receptor_binding.html for an interactive heatmap that shows effects of mutations across the full spike, as well as interactive versions of other panels of this figure. **B**. Correlation between ACE2 binding measurements for the two independent deep mutational scanning library replicates faceted by proximity to ACE2. ACE2 proximal sites are defined as those within 15 Å distance from ACE2 in ACE2-bound RBD structure (PDB: 6M0J). **C**. Correlation between the effects of RBD mutations on ACE2 binding measured for the KP.3.1.1 spike in the current study and the XBB.1.5 spike in prior work (2). **D**. Mutation effects on ACE2 binding at sites 435 and 493 measured in XBB.1.5 versus KP.3.1.1 deep mutational scanning libraries. Amino-acids are coloured by their chemical properties.

There is a good correlation between the effects of RBD mutations on ACE2 binding in our KP.3.1.1 deep mutational scanning and similar data previously published for the XBB.1.5 spike (2) (**Fig. 3C**). However, there are some mutations with different effects on ACE2 binding in KP.3.1.1 and XBB.1.5, including A435S which increases binding to ACE2 in KP.3.1.1 but decreases binding for XBB.1.5 (note this mutation also had contrasting effects on cell entry in the two spikes as described above) (**Fig. 3D**). The A435S mutation has been rare for most of SARS-CoV-2’s evolution, but has recently occurred independently in multiple lineages including the JN.1-descendants NB.1.8.1, XEC.25, MC.10.1, MC.31, and NP.1 variants and a recent BA.3-descendant saltation variant BA.3.2. In addition, E493D and E493N increase ACE2 binding by the KP.3.1.1 spike, but in XBB.1.5 mutating site 493 from its initial identity of Q to any of E, D, or N impairs ACE2 binding (**Fig. 3D**) (2,34).

### Mutation effects on serum neutralization

We measured how spike mutations affect neutralization by sera collected from seven human individuals pre- and post-exposure by vaccination or infection with JN.1-descendant variants (**Table S1**). All seven individuals were adults who had originally been imprinted by vaccination with the early COVID-19 vaccine in 2021 followed by various further booster vaccinations and infections. For most (although not all) of these individuals, exposure to a JN.1-descendant spike via vaccination increased neutralizing serum titers against KP.3.1.1 (**Fig. S3A-B**).

We used the pseudovirus libraries to measure how the KP.3.1.1 spike mutations affected neutralization by the sera from each individual both pre- and post-vaccination or infection with a JN-1 descendant spike. For the most part, mutations had similar effects on neutralization by sera from each individual collected pre-versus post-vaccination or infection (**Fig. 4** and **S3C**). Across all sera, the sites where mutations caused the most escape from serum neutralization were primarily in the RBD (**Fig. 4** and interactive plot at https://dms-vep.org/SARS-CoV-2_KP.3.1.1_spike_DMS/polyclonal_sera_escape.html). RBD mutations at sites 332, 344, 357, 393, 428, 458, 470 and 518 caused the greatest serum escape both pre- and post-vaccination or infection (**Fig. 4A**). Some sites outside the RBD also reduced serum neutralization, including sites 50, 132, 200, 222 in NTD, 572 in SD1, and 852 in S2. Notably, most of the sites where mutations caused the greatest escape in the RBD and all the strongest sites of escape outside the RBD are ones where mutations affect ACE2 binding (**Fig. 3B**, and next section), suggesting mutations at these sites impact serum neutralization largely changing the RBD’s up/down conformation, thereby indirectly affecting binding by antibodies targeting potent neutralizing epitopes on the RBD (2,35–37). However, there are also some sites of appreciable escape where mutations do not affect RBD up/down binding (e.g., 456, 458, 475, 478, 487); these mutations likely directly escape binding by neutralizing antibodies rather than affecting RBD up/down conformation.

**Figure 4.**
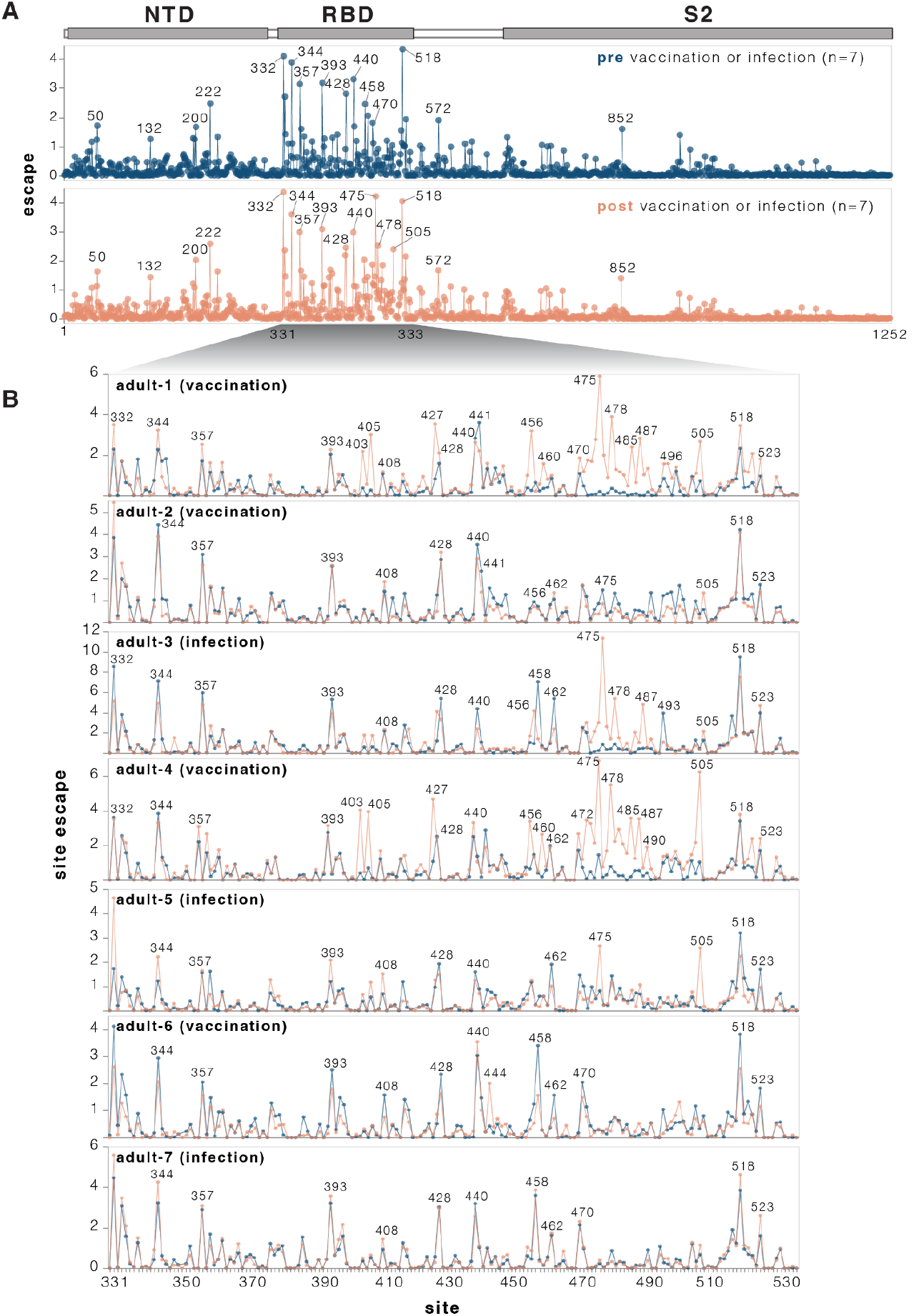
Effects of mutations to the KP.3.1.1 spike on serum neutralization. **A**. Total neutralization escape by all measured mutations at each site in spike averaged across all seven pre- or post-vaccination or infection sera. For more extensive interactive versions of the plots in this figure including heatmaps with per-mutation effects, see https://dms-vep.org/SARS-CoV-2_KP.3.1.1_spike_DMS/polyclonal_sera_escape.html. **B**. Comparison between escape at RBD sites pre- and post-vaccination or infection for each of the seven individual sera. Note that this plot only shows positive escape values (mutations that reduce neutralization), and not mutations that increase neutralization (negative escape), although the interactive plots linked in this legend have options to view the negative escape.

While many mutations that reduce serum neutralization pre- and post-vaccination or infection were shared among the different sera, in a subset of individuals exposure to a JN.1-descendant spike clearly shifts neutralization immunodominance. In adult-1, adult-3, and adult-4 and adult-5, several RBD sites where mutations had little or no effect on serum neutralization before JN.1-descendant spike exposure become the dominant escape sites after vaccination or infection (**Fig. 4B**). These new escape sites include 403, 405, 475, 478, 487, 490 and 505. Notably, in circulating SARS-CoV-2 variants, many of these sites have recently acquired mutations that reduce serum neutralization. For example, the XFJ, JN.1.18.5, LF.7.1.2, LF.7.2.1, PC.2 and LP.8.1.9 variants all carry A475V, BA.3.2 carries K478N while NB.1.8.1 carries K478I, and XFG carries N487D.

We validated the deep mutational scanning measurements of how mutations affect serum neutralization using standard SARS-CoV-2 pseudovirus neutralization assays (**Fig. S4**) (38). The deep mutational scanning measurements correlated well with changes in IC50 values measured in the standard neutralization assays (**Fig. S4A**). We also confirmed via standard neutralization assays that mutations A475V, H505E, K478I and N487D cause a larger reduction in the neutralization by the serum from some individuals after versus before exposure to a JN.1-descendant spike (**Fig. S4B**), consistent with the deep mutational scanning.

### Sites where mutations affect RBD up/down conformation

To identify sites in spike that affect RBD up/down conformation, we leveraged the previously noted fact that mutations at these sites have opposing effects on ACE2 binding and serum antibody neutralization escape: namely, mutations that put the RBD more in the up conformation increase ACE2 binding but also enhance neutralization (2,35–37). Our measurements for the KP.3.1.1 spike show this relationship clearly: there is a strong inverse correlation between serum neutralization escape and ACE2 binding for mutations that affect both these phenotypes but are distal from the RBD’s ACE2-binding motif (**Fig. 5A**). This inverse correlation is due to the fact that positioning RBD in the up conformation reveals the receptor-binding motif, which mediates binding to ACE2 but is also targeted by many potent neutralizing antibodies. Therefore, mutations that put the RBD more in the up conformation sensitize the spike to serum neutralization (negative escape values in our measurements), while mutations that put the RBD more in the down conformation tend to cause serum neutralization escape. By contrast, ACE2-proximal sites show no correlation between ACE2 binding and serum neutralization (**Fig. 5A**) because they often both interact with the receptor directly and are directly targeted by neutralizing serum antibodies. Note, that some ACE2 proximal sites may still modulate the RBD up/down conformation, but this modulation does not lead to the aforementioned consistent pattern on ACE2 binding and neutralization because the direct effects of mutations at these sites both ACE2 binding and neutralizing antibody binding can overwhelm the effect of the RBD up/down conformation modulation.

**Figure 5.**
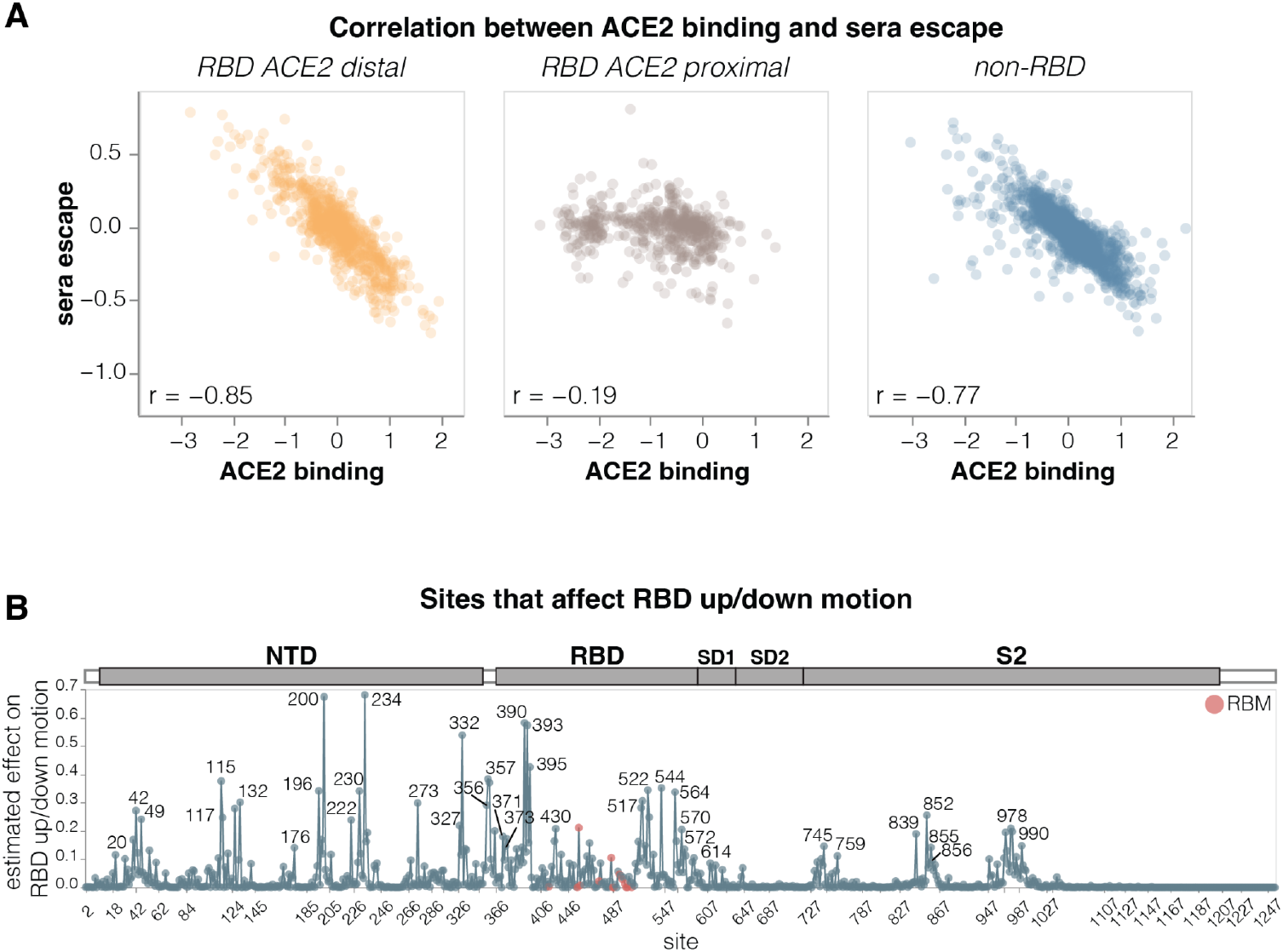
Sites where mutations affect RBD up/down conformation. **A**. Correlation between the measured effects of each mutation on ACE2 binding and serum antibody escape, faceted by proximity of the site to ACE2. **B**. Experimentally estimated effect of mutations at each site on RBD up/down conformation. The larger the value, the greater effect mutations at that site have on RBD up/down conformation, although individual mutations at each site may have opposing effects. Sites within the receptor-binding motif (RBM) are colored red, and all other sites are blue. See https://dms-vep.org/SARS-CoV-2_KP.3.1.1_spike_DMS/RBD_movement.html for an interactive version of this plot. The effect of each site on RBD up/down conformation is estimated from the deep mutational scanning by calculating correlation (Pearson R) between serum neutralization escape and ACE2 binding for all mutations at each site, then multiplying that correlation by minus one and weighting it by the root-mean-square (RMS) effect of all mutations at the site on ACE2 binding and the RMS effect of all mutations at the site on serum neutralization escape. Sites with positive correlation had the effect floored to zero. This metric captures the fact that mutations at sites that affect RBD up/down conformation have opposing effects on ACE2 binding and serum neutralization escape. Only sites where binding and neutralization effects could be measured for at least three mutations are shown.

To estimate how much each site affects RBD up/down conformation, we calculated the correlation between serum neutralization escape and ACE2 binding at each site, weighting it by the root mean square effect of mutations at each site on both phenotypes (**Fig. 5B** and interactive plot at https://dms-vep.org/SARS-CoV-2_KP.3.1.1_spike_DMS/RBD_movement.html). Among the sites that stand out as strongly affecting RBD up/down conformation are many clade-defining mutations as well as some of the most frequently mutated sites through various periods of SARS-CoV-2 evolution in humans. Site 222 was one of the most frequently mutated sites just before Omicron emerged (39), sites 371 and 373 fixed mutations in all Omicron lineages (40), and sites 332, 356 and 570 fixed mutations in the BA.2.86-lineage which is the ancestor of most currently circulating strains (41). The prevalence of mutations at sites that modulate RBD up/down conformation in major SARS-CoV-2 lineages suggests a strong selective pressure to balance receptor binding with resistance to neutralization by RBD-directed antibodies; indeed evidence suggest that multiple recent SARS-CoV-2 variants have acquired mutations that position the RBD in a more closed conformation (36,42).

### Effects of mutations on neutralization by clinically relevant monoclonal antibodies

We next determined how mutations to spike affect neutralization by three clinically relevant monoclonal antibodies: BD55-1205 (12), SA55 (43), and VYD222 (44,45) (**Fig. 6**). BD55-1205 and SA55 have maintained high neutralizing potency against currently circulating variants (12). SA55 is in clinical trials in China (41), BD55-1205 is licensed to Moderna Inc. (12), and VYD222 is currently the only SARS-CoV-2 antibody authorized for use in the USA for pre-exposure prophylaxis in immunocompromised individuals (it is the antibody in Pemivibart) (47). Knowledge of which mutations reduce neutralization by these antibodies is important for ongoing surveillance, as all other clinically approved SARS-CoV-2 antibodies have now been escaped by viral mutations (43,48).

**Figure 6.**
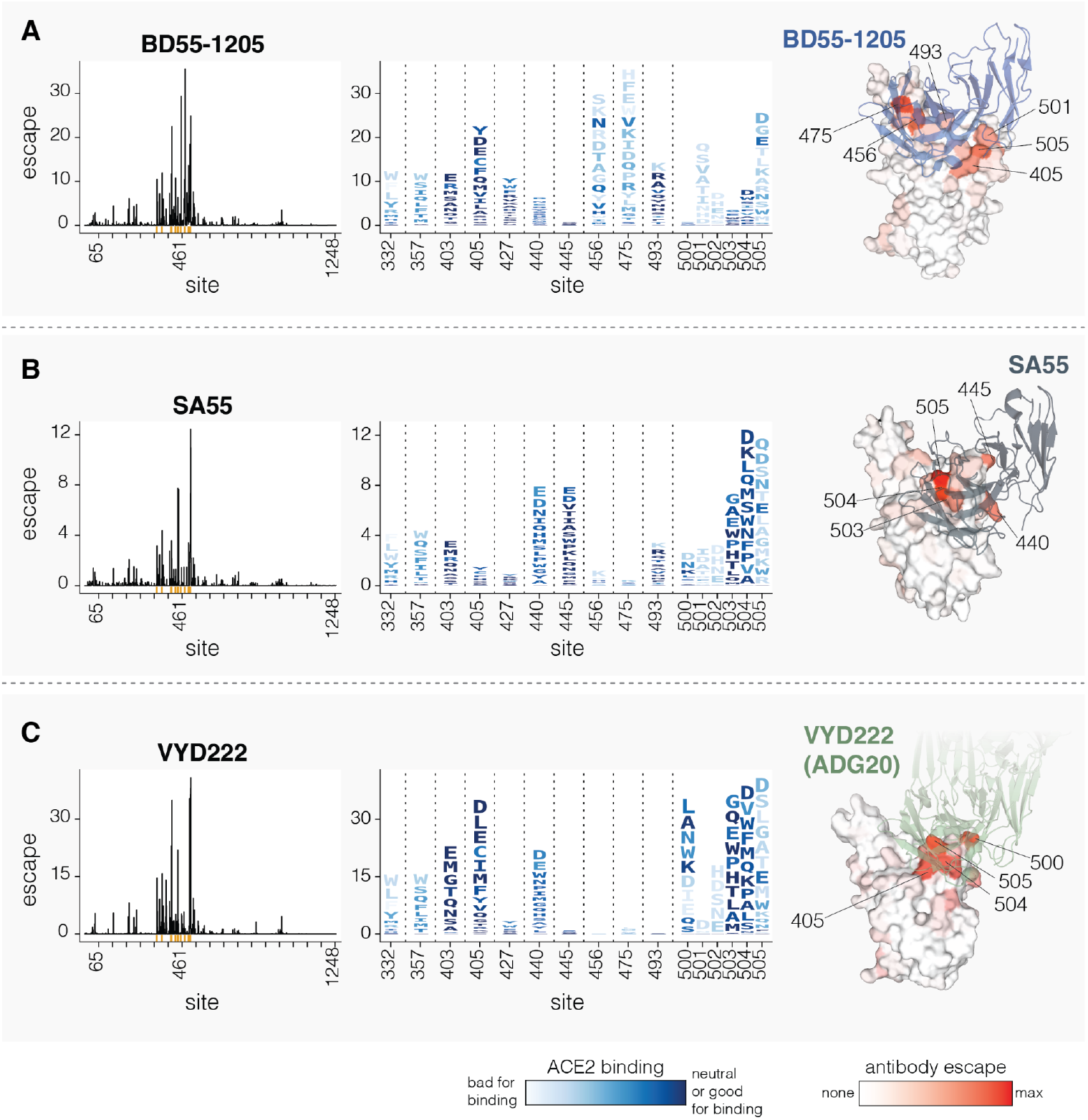
Mutations that reduce neutralization by monoclonal antibodies BD55-1205, SA55 and VYD222. **A**. Mutations that reduce neutralization by the BD55-1205 antibody. The line plot on the left shows the total escape caused by all mutations at each site in spike. The logo plot in the middle shows escape caused by each mutation at key sites; letter heights indicate escape caused by each mutation, and mutations are colored by their effect on ACE2 binding. The structure at right shows a surface representation of the RBD bound by BD55-1205, with the RBD colored by the total escape at each site (PDB ID: 8XE9). **B-C**. Same as A but for SA55 and VYD222, respectively. For SA55, the structure is PDB ID 7Y0W For VYD222, the structure is PDB ID 7U2D, which shows ADG20, which is the parent antibody from which VYD222 is derived (47). Only positive escape values (mutations that reduce neutralization) are shown. For a more detailed interactive plot showing mutation-level escape across the spike for all three antibodies, see https://dms-vep.org/SARS-CoV-2_KP.3.1.1_spike_DMS/antibody_escape.html.

All three antibodies bind to sites around the RBD’s receptor binding motif, with SA55 and VYD222 sharing especially similar structural epitopes (43,49). Our deep mutational scanning shows that all three antibodies are strongly affected by mutations at several sites in the range from 500 to 505, although the exact sites in this range where mutations have the most impact varies among the antibodies (**Fig. 6**). BD55-1205 neutralization is also affected by mutations at sites 456, 475 and 493, all of which interact with ACE2 (**Fig. 6A**). Most changes to 456 and 475 sites are deleterious for ACE2 binding (see letter colors in logoplots in **Fig 6**), although A475V, which measurably escapes BD55-1205, is only mildly deleterious for ACE2 binding and has recently occurred in several JN.1-descendant lineages. SA55 is affected by mutations to sites 440 and 445, and to a lesser degree by mutations at 493 (**Fig. 6B**). VYD222 is also affected by mutations at site 440 in addition to mutations at sites 405 and 403 (**Fig. 6C**). Because sites 500-505 are primarily accessible in the RBD’s up position, all three antibodies are affected by mutations that modulate RBD up/down movement, as has been noted previously (12,50,51). In particular, some mutations at sites 332, 357, and 427 affect neutralization by all three antibodies to various degrees, despite the fact none of these sites are in the direct structural epitopes, presumably by putting the RBD more in the down conformation and so partially shielding the antibody epitopes.

Interestingly, in our pseudovirus deep mutational scanning, mutations at site 505 cause significantly more escape from all three of the antibodies than reported in previously published yeast-based RBD-only deep mutational scanning data suggest (**Fig. S5**) (6,43,45). We hypothesize that this difference is because RBD-only assays measure just the direct effects of mutations on antibody-RBD binding, whereas the pseudovirus deep mutational scanning also measures the impacts of mutations on RBD up/down movement that affect RBD epitope accessibility in the context of full spike. Indeed, mutations at RBD motion-regulating sites 332, 357 and 427 affect neutralization by all three antibodies in full-spike but not in yeast-based RBD-only deep mutational scanning (**Fig. S5**). Similarly, mutations at site 505 both directly affect antibody-RBD binding and the up/down motion of the RBD due to this site’s location in the inter-protomer interface in the down RBD spike conformation. While site 505 is likely under significant evolutionary constraint because most mutations at that site reduce ACE2 binding, our serum-escape measurements described above suggest this site may be starting to come under appreciable pressure for mutations from population immunity.

## Discussion

Here we have measured how mutations to the KP.3.1.1 spike affect several distinct phenotypes: cell entry, ACE2 binding, serum neutralization, RBD up/down motion, and neutralization by key monoclonal antibodies. These measurements provide several important insights into the selection pressures and molecular constraints currently shaping SARS-CoV-2 evolution.

First, our measurements underscore the substantial impact of mutations that affect RBD up/down motion on receptor binding and antibody neutralization. In the context of the full spike, mutations that affect RBD up/down motion impact ACE2 binding as much as mutations at RBD sites that interact with ACE2 directly. Mutations that affect RBD up/down motion have a consistent signature: they have opposite effects on ACE2 binding and serum neutralization, since putting the RBD more up increases the accessibility of the receptor-binding motif to bind ACE2 but also makes it more susceptible to RBD-targeting neutralizing antibodies (33,52). Many sites that affect RBD up/down motion have mutated in major lineages during the course of SARS-CoV-2 evolution in humans, emphasizing the importance of the balancing effects of RBD up/down movement on viral fitness via impacts on ACE2 binding and serum neutralization. Note that mutations that put the RBD in a more up conformation may promote the cross-species transfer of coronaviruses by increasing binding to receptors from new species (32,53,54); it appears that the spike of the first SARS-CoV-2 strains identified in humans had the RBD in a relatively more up conformation, and subsequent evolution has selected for mutations that position the RBD more down (36,42).

Second, our measurements identify sites where mutations cause the largest reductions in neutralization by human serum antibodies; there are already newly emerging viral lineages that carry some of these mutations. Most of the sites where mutations most impact serum neutralization are in the RBD as expected from prior work showing that RBD-directed antibodies are usually responsible for most serum neutralizing activity (55–57), although mutations at some NTD sites also have a substantial effect. Some of the top RBD sites of serum antibody escape are likely directly in the epitopes of neutralizing antibodies that sterically block receptor binding (e.g., sites 456, 458, 475, 478, 487); mutations at some of these sites have recently been observed in new SARS-CoV-2 lineages. However, mutations at NTD and RBD sites that affect RBD up/down motion and so affect serum neutralization indirectly by conformational masking, often have as much impact on serum neutralization as direct escape mutations in key RBD epitopes. As mentioned above, some of these up/down affecting sites have mutated in major lineages; however, such conformational escape is constrained by the fact that mutations that reduce serum neutralization by putting the RBD in a more down conformation also reduce ACE2 binding, and so may need to be buffered by other ACE2 affinity increasing mutations.

Third, we find that exposure to a JN.1-descendant spike (via vaccination or infection) often shifts the neutralization immunodominance hierarchy to new epitopes. Specifically, for some individuals, vaccination or infection with a JN.1-descendant variant leads to mutations at new sites causing large reductions in neutralization; these new sites include several (e.g., 475, 478, and 487) that have acquired mutations in very recent SARS-CoV-2 lineages. Our data cannot determine the underlying mechanism responsible for this shift in serum neutralizing specificity. Once individuals have been imprinted by SARS-CoV-2 infection or vaccination, most of the neutralizing response to subsequent vaccinations and infections is driven by activation of pre-existing cross-reactive B cells (28,58–61). However, the affinity maturation of these pre-existing B cells can shift the balance of epitope targeting in polyclonal sera (59). In addition, sufficient exposures to new variants can activate naive B cells (61,62). The shifts in serum neutralizing specificity we observe after exposure to a JN.1-descendant variation could be due to some combination of boosting of pre-existing cross-reactive B-cells that were previously subdominant, affinity maturation of pre-existing B-cells to better target recently mutated epitopes, or activation of naive B-cells targeting new epitopes. Regardless of the underlying mechanism, the fact that exposure to recent variants changes the neutralization immunodominance hierarchy supports the idea that updating vaccines to more recently circulating variants (23) can shift the specificity of neutralizing antibodies to target newer SARS-CoV-2 variants.

The fact that exposure to recent JN.1-descendant variants can shift which spike mutations affect neutralization by the serum antibodies of imprinted adults highlights the increasing heterogeneity in antibody immunity across the human population. We recently showed that the epitopes targeted by the neutralizing antibodies of young children who had experienced just a single infection with a recent variant differ dramatically from those targeted by adults imprinted by infection or vaccination early in the SARS-CoV-2 pandemic (20). The current study only examined serum from imprinted adults, but finds heterogeneity even among such adults depending on whether they have been exposed to a JN.1-descendant variant. This increasing immune heterogeneity across the population may favor more co-circulation of multiple SARS-CoV-2 lineages rather than repeated rapid evolutionary sweeps by a single variant (63,64).

We also mapped how mutations affect neutralization by three clinically relevant monoclonal antibodies (BD55-1205, SA55 and VYD222) that have so far retained neutralizing activity against nearly all SARS-CoV-2 lineages (65,66). A major epitope targeted by all these antibodies is the 500-505 loop in the RBD, which has not mutated in any major lineage since the emergence of Omicron in 2021. Notably, these antibodies target functionally constrained RBD epitopes that overlap with the ACE2 binding motif and are only fully accessible in the up RBD conformation, and our data show that neutralization by all three antibodies is reduced by mutations that put the RBD in a more down conformation. In particular, mutations to site 505, which both affects RBD motion and forms part of the epitope for all three antibodies, have a greater impact on pseudovirus neutralization than was apparent in prior RBD-only yeast-display deep mutational scanning (6). Site 505 remains under substantial constraint, since most mutations at that site both reduce direct RBD-ACE2 binding affinity (15) and put the RBD in a more up conformation that increases its susceptibility to RBD-directed serum neutralizing antibodies. However, our results show that site 505 is now a serum neutralization escape mutation for some individuals who have been exposed to a JN.1-descendant variant, suggesting such individuals now produce appreciable neutralizing antibodies directly targeting site 505. Therefore, site 505 might be under increasing pressure to mutate in circulating SARS-CoV-2 lineages, although additional changes to spike would likely be needed to overcome the pleiotropic effects such a mutation would have on ACE2 binding and RBD up/down conformation.

## Acknowledgements

We thank David Veesler from University of Washington for providing soluble ACE2 protein. We thank Ryan Hisner and Federico Gueli for useful comments on the manuscript. This research was funded by grants from the NIAID/NIH awarded to JDB: P01AI167966 and the SAVES program (contract 75N93021C00015, option 18.C). JDB is an investigator at the Howard Hughes Medical Institute. This research was also supported by the Genomics & Bioinformatics Shared Resource, RRID:SCR_022606, of the Fred Hutch/University of Washington Cancer Consortium (P30 CA015704), by the Flow Cytometry Shared Resource, RRID:SCR_022613, of the Fred Hutch/University of Washington/Seattle Children’s Cancer Consortium (P30 CA015704), and by Fred Hutch Scientific Computing, NIH grants S10-OD-020069 and S10-OD-028685. SH is a postdoctoral fellow of the Translational Data Science Integrated Research Center at the Fred Hutchinson Cancer Center. BBL is a Washington Research Foundation postdoctoral fellow. This material is based upon work supported by the National Science Foundation Graduate Research Fellowship under Grant No. DGE-2140004. Any opinion, findings, and conclusions or recommendations expressed in this material are those of the authors and do not necessarily reflect the views of the National Science Foundation. This manuscript is the result of funding in whole or in part by the National Institutes of Health (NIH). It is subject to the NIH Public Access Policy. Through acceptance of this federal funding, NIH has been given a right to make this manuscript publicly available in PubMed Central upon the Official Date of Publication, as defined by NIH.

## Methods

### Data availability and interactive figures

All data described in this manuscript are available as raw numerical values and in various interactive figure formats:

- Interactive figures can be found at a website associated with this manuscript https://dms-vep.org/SARS-CoV-2_KP.3.1.1_spike_DMS/. The website homepage has interactive figures organised by phenotype and by clicking on each phenotype the reader can find key plots, descriptions and links to raw numerical values used to make the interactive plots.
- The computational analysis pipeline used to analyse deep mutational scanning data and make all associated manuscript figures is on GitHub at https://github.com/dms-vep/SARS-CoV-2_KP.3.1.1_spike_DMS.

Sequencing data associated with this manuscript has been deposited to the SRA under BioProject PRJNA1305008..

### Deep mutational scanning library design

Deep mutational scanning libraries were designed to cover all possible mutations in the RBD and all tolerated and frequently mutated changes outside the RBD. To identify tolerated and frequently mutated sites we included mutations that occur more than 50 times among SARS-CoV-2 genomes deposited on GISAID (67), mutations that occur at least 10 times on UShER (68) spike phylogenetic tree, any mutation present in a recent SARS-CoV-2 lineage (at the time of library design these lineages where BA.2.86, JN.1, JN.1.11.1, and KP.3), and any mutations that occurred at least once in a Pango designated lineage (69). In addition, we introduced all possible amino-acid mutations at sites that fit any of the following criteria: mutated at least 50 times in a recent SARS-CoV-2 lineage, mutated along UShER spike phylogenetic tree at least 2500 times, mutated repeatedly at least 3 times among any Pango designated lineages, or had mutated in KP.3.1.1 variant relative to Wuhan-Hu-1 sequence. The above criteria were also applied for deletions but deletions were only included if they were present at any site in the NTD or positions 331-354 or 434-508 in the RBD. Several mutations and sites to saturate were also included manually in library design regardless of their frequency counts based on reports of these mutations occurring in circulating lineages at the time of library design. The list of manually included mutations as well as parameters for all other selection criteria is at https://github.com/dms-vep/SARS-CoV-2_KP.3.1.1_spike_DMS/blob/main/library_design/config.yaml. The full list of all mutations included in the library design is at https://github.com/dms-vep/SARS-CoV-2_KP.3.1.1_spike_DMS/blob/main/library_design/results/mutations_to_make.csv.

### Overview of library construction using Golden Gate assembly

Golden Gate assembly was used to create KP.3.1.1 spike coding plasmid libraries containing all the designed mutations (70–76) (**Fig. S6**). Due to the length of the spike sequence and the number of mutations we wanted to include in the library it was cost-prohibitive to synthesize the spike gene as a single fragment for all spike variants we wanted to include. We therefore subdivided spike into 17 overlapping tiles between 250-290 nt in length (close to the maximum length that can be synthesized by Twist Bioscience as a single-stranded DNA (ssDNA) oligo pool) (**Fig. S6A**), computationally designed a pool of oligos, where each oligo is one of the tiles with a mutation we wanted to include in the library (**Fig. S6B**), and ordered all the oligos pooled together as ssDNA fragments from Twist Bioscience. From that ssDNA pool we performed 17 individual PCR reactions to amplify oligos belonging to each tile using primers with flanking sequences containing BsmBI restriction sites (**Fig. S6C**). Golden Gate assembly was then used to assemble each tile pool and flanking spike sequences unique to each tile into a shuttle vector (**Fig. S6D**). The assembled shuttle vector pool was electroporated into bacteria and next day plasmids were recovered for all 17 pools. The full spike sequence was amplified from each pool using primers with flanking sequences that match lentiviral backbone as well as a barcode sequence in the reverse primer (**Fig. S6E**). All 17 barcoded spike pools were then pooled equimolarly and HiFi assembly was used to clone the library (**Fig. S6F**), which after pooling had all designed mutations throughout the spike, into a lentivirus backbone.

The sequence of the codon optimized KP.3.1.1 spike in the final lentiviral backbone used to make pseudovirus-based libraries is at https://github.com/dms-vep/SARS-CoV-2_KP.3.1.1_spike_DMS/blob/main/library_design/data/4838_pH2rU3_ForInd_KP.3.1.1_sinobiological_CMV_ZsGT2APurR.gb. Sequences for all 17 overlapping tiles are at https://github.com/dms-vep/SARS-CoV-2_KP.3.1.1_spike_DMS/blob/main/library_design/data/KP311_GAA_assembly_fragments.csv. Tiles were designed manually making sure that the overhangs for the fragments that will be assembled during the Golden Gate assembly step are unique for each fragment and have a sequence compatible with high fidelity assembly (77). The 1st and the 17th tile overlapped with a pGGAselect DNA shuttle vector that is provided in NEBridge® Golden Gate Assembly Kit (BsmBI-v2) (E1602L). The oligo pool was designed using a script available at https://github.com/jbloomlab/gga_codon_muts_oligo_design. The script reads in tile sequences and desired mutation spreadsheet and generates a fasta file with oligo sequences that can be uploaded directly for ordering oligo pool from Twist Biosciences. We set the oligo design script to intentionally include 0.005 fraction of unmutated sequences for each tile in order to have some wild-type KP.3.1.1 spike in the final pseudovirus library, as well as avoid any mutation design that would introduce BsmBI cut sites. Sequences for designed oligos covering all 17 tiles is at https://github.com/dms-vep/SARS-CoV-2_KP.3.1.1_spike_DMS/blob/main/library_design/results/mutagenesis_oligos.fa. A GitHub repository that selects the mutations to be included in the library and designs mutated oligos for each tile is at https://github.com/dms-vep/SARS-CoV-2_KP.3.1.1_spike_DMS/tree/main/library_design.

### Deep mutational scanning plasmid library cloning using Golden Gate Assembly

To amplify individual tile pools from one ssDNA oligo pool we performed 17 PCR reactions. For each reaction we used KOD Hot Start Master Mix (Sigma-Aldrich, Cat. No. 71842), 0.3 µM of forward and reverse primer and 2 ng of ssDNA oligo pool. Each reaction was started at 95°C for 2 min and then went through 23 cycles of 95°C for 20 s, 62°C for 10 s, 68°C for 25 s. To amplify flanking spike sequences for each tile we used KOD Hot Start Master Mix, 0.3 µM of forward and reverse primer and 1 ng of KP.3.1.1 spike coding lentiviral backbone (see above section for plasmid map). The full list for forward and reverse primers used in both reactions is at https://github.com/dms-vep/SARS-CoV-2_KP.3.1.1_spike_DMS/blob/main/library_design/data/primers.csv. Expected size products were gel and Ampure XP bead purified (1:3 DNA to bead).

We then performed Golden Gate assembly using NEBridge Golden Gate Assembly Kit (BsmBI-V2). For the assembly we used 100 fmol of amplified tile pool and flanking spike sequence fragments each and 50 fmol of pGGAselect shuttle plasmid (provided in NEBridge Golden Gate Assembly Kit). The assembly reactions were incubated at 42°C for 1 min followed by 16°C for 1 min for 30 cycles, followed by 60°C for 5 min. The reactions were then purified using Ampure XP beads and eluted in 20 µl of water. 1 µl of purified assembly was then used to electroporate NEB® 10-beta Electrocompetent E. coli cells (C3020K). Electroporated cells were then suspended in 1 ml of recovery media and shaken at 37°C for 1 hour. After recovery, cells were spun down, recovery media was removed and cells were resuspended in chloramphenicol-containing LB media for incubation at 37°C with shaking overnight. High transformation efficiency (∼1 million colonies per tile library) was confirmed by diluting a small amount of recovered cells, plating on chloramphenicol-containing agar plates overnight and counting colony forming units the next day. High transformation efficiency at this and later steps is important to avoid any barcode duplication at later virus production steps due to lentivirus recombination. Note also that here and in later electroporation steps we used liquid cultures to amplify our plasmid libraries as opposed to high-density spread on bacterial culture plates we used in the past as this has been shown to be sufficient for a uniform plasmid amplification (78). After overnight growth, shuttle plasmid libraries for each tile were recovered using QIAprep Spin Miniprep Kit (Cat. No. 27106X4).

Next, the spike libraries for each tile were amplified and barcoded. We performed PCR on each tile plasmid library using KOD Hot Start Master Mix, 10 ng of plasmid library and 0.3 µM of forward (5′-gcacgcgCAGCCGAGCCACATCGCTCA-3′) and reverse (5′-gcggaactccactaggaacatttctctctcgaaTCTAGANNNNNNNNNNNNNNNNAGATCGGAAGAGCGTCGTGTAGGGAAAGAG-3′) primers, the latter primer contained a 16 nt barcode. After amplification each spike tile library was purified by gel and Ampure XP beads. Note gel purification at this step is important because we found cloning of some tiles produces a minor amount of truncated spike and gel purification allowed us to recover only the full length products. All barcoded spike libraries were then pooled equimolarly. We made two equimolar pools of barcoded spike libraries to make library-1 and library-2 biological replicates. All subsequent steps in library production were done in parallel for library-1 and library-2. NEBuilder® HiFi DNA Assembly Master Mix (E2621S) was then used to assemble barcoded spikes into a lentivirus backbone, as described previously (21). See lentivirus backbone structure in **Fig. S6F**; plasmid for the backbone is available at Addgene #204579). Assembled backbones were electroporated in electrocompetent bacteria and plasmids were amplified using liquid culture,as described above. As before we confirmed high electroporation efficiency at this step and cultured at least 10 million colony forming units per library replicate.

### Production of cell-stored deep mutational scanning libraries

To produce the cell-stored deep mutational scanning libraries we used a method described previously (**Fig. S1A**) (21). In brief, we first used lentivirus backbones that carried barcoded spike libraries to produce VSV-G pseudotyped viruses. To do so we transfected two 6-well plates of 293T cells with lentivirus helper plasmids (BEI: NR-52517, NR-52519, NR-52518) and VSV-G expression plasmid (Addgene #204156). 48 hours after transfection we collected VSV-G pseudotyped viruses from cell supernatant and used them to infect 293T-rtTA cells at low multiplicity of infection (<0.01) so that most infected cells were infected with only one viral variant. We then used puromycin to select for successfully transduced cells. The transduced cell library pool was then expanded and frozen at >15 M cells per aliquot in liquid nitrogen until further use.

### Long-read sequencing for variant-barcode linkage

To build a variant to barcode lookup table for the deep mutational scanning libraries, we rescued VSV-G pseudotyped viruses from the cell-stored libraries. We use VSV-G pseudotyping at this stage to rescue all virus variants from the cells regardless of how deleterious a mutation in spike may be. To do so we transfected library cells with lentivirus helper plasmids and VSV-G expression plasmid and 48 hours after transfection we recovered VSV-G pseudoviruses from cell supernatant, purified them from cell debris using 0.45 µm SFCA Nalgene 500mL Rapid-Flow filter unit (Cat. No. 09-740-44B), and concentrated using Pierce Protein Concentrator (ThermoFisher, 88537). We then used ∼10 million transcription units of VSV-G pseudotyped viruses to infect 293T cells and 15 hours after infection recovered non-integrated viral genomes using QIAprep Spin Miniprep Kit. We then performed two rounds of PCR to amplify the barcoded spikes in the recovered lentivirus genomes, minimizing the number of PCR cycles to avoid strand-switching. Long-read circular consensus sequencing was performed on amplified virus genomes using PacBio Sequel IIe machine. Consensus sequence for each variant was determined using at least 2 CCSs per barcode. Variant-barcode lookup table for both biological KP.3.1.1 library replicates is at https://github.com/dms-vep/SARS-CoV-2_KP.3.1.1_spike_DMS/blob/main/results/variants/codon_variants.csv. On average each variant had 1.25 and 1.27 mutations per spike for library-1 and library-2, respectively.

### Measurement of mutation effects on cell entry effect

KP.3.1.1 spike pseudotyped viruses were produced from cell-stored libraries as described previously (2). 150 million library cells were plated into 5-layer flasks (Corning Falcon 875cm^2^ Rectangular Straight Neck Cell Culture Multi-Flask, Cat. No. 353144) in the presence of 1 µg/ml of doxycycline to induce spike expression from TRE3G promoter in the lentivirus backbone. Next day cells were transfected with 50 µg of each lentiviral helper plasmid and during transfection cell media was replaced with fresh serum-free media (Opti-MEM supplemented with 0.1% heat-inactivated FBS, 0.3% bovine serum albumin, 100 µg/mL of calcium chloride, 100 U/mL penicillin, and 100 µg/mL streptomycin). Serum free media was used because it allows better virus concentration in protein columns as FBS tends to clog column filters. 48 hours after transfection cell supernatant was collected, purified from cell debris and concentrated using protein columns. Protein column concentrated virus titers varied between 12-25 million transcription units per ml. VSV-G pseudotyped viruses were also produced in parallel to spike pseudotyped libraries, using protocol described in the section above. For cell entry effect measurements both 3 million transcription units of spike pseudotyped libraries and 10 million transcription units VSV-G pseudotyped libraries were used to infect medium-ACE2 (24) cells and 293T cells, respectively. For spike pseudotyped library infections cells were plated in the presence of 2.5 µg/ml of amphotericin B (Sigma, Cat. No. A2942), which we have shown in the past increases virus titers (21). 15 hours after infection non-integrated viral genomes were recovered using QIAprep Spin Miniprep Kit and amplicon libraries were prepared for illumina sequencing as described previously using dual indexing for each sample to avoid index hopping on certain sequencing platforms (21). Sequencing was performed on NovaSeq X Plus and NextSeq 2000 platforms.

Mutation effects on cell entry were calculated using log enrichment ratio: 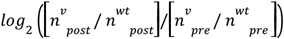, where 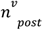 is variant count post-infection (spike pseudotyped virus infection), 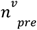 is variant count pre infection (VSV-G pseudotyped virus infection) and 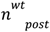 and 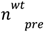 are unmutated variant counts post- and pre-infection. The *multidms* (79) package was used to fit global epistasis models (80) on variant effect data to estimate the effects of individual mutations from the full libraries of both singly and multiply mutated spike variants. The values reported here are the median across the measurements with all replicates of both libraries.

### Measurement of mutation effects on receptor binding

To measure how mutations to spike affect ACE2 binding we used soluble monomeric ACE2. Monomeric ACE2 was produced as described previously (2). First, we mixed 1.5 million transcription units of spike pseudotyped library virus per sample with RDPro pseudotyped virus at 1-2 % of total transcription units used. Use and production of RDPro pseudotyped virus was described previously (2). RDPro is used in our experiments as a non-neutralizable standard to convert sequencing counts to fractional neutralization of each variant at each ACE2 concentration as described previously (2). The library virus was then mixed with increasing concentrations of soluble monomeric ACE2 and incubated at 37°C for 30 min. The ACE2 concentrations were selected such that they would cover most of the KP.3.1.1 spike pseudotyped virus neutralization range in order to identify mutations that both increase (spike variants that are neutralized well at low ACE2 concentrations) and decrease (spike variants that are neutralized at high ACE2 concentrations) ACE2 binding; specific concentrations used were 6, 13, 27, 54, and 115 µg/ml. After incubation, libraries were used to infect medium-ACE2 cells in the presence of 2.5 µg/ml of amphotericin B and 15 h post infection non-integrated viral genomes were recovered and prepared for Illumina sequencing as described previously (21). After converting the sequencing counts to the fractional neutralization using the non-neutralized RDPro standard (2), we analyzed the data using a biophysical model implemented in the *polyclonal* software package (https://github.com/jbloomlab/polyclonal) (81) to determine the effect of each mutation on ACE2 neutralization, reporting the values such that positive effects indicate improved ACE2 binding (higher neutralization by soluble ACE2). We performed ACE2 binding experiments with both library-1 and library-2 biological replicates. The values reported here are the median across both replicates. Mutations effects on ACE2 binding are shown at https://dms-vep.org/SARS-CoV-2_KP.3.1.1_spike_DMS/receptor_binding.html.

### Measurement of mutation effects on serum and antibody neutralization

Before performing sera and antibody selection experiments with deep mutational scanning libraries we determined their potency by performing pseudovirus neutralization assays on viruses pseudotyped with KP.3.1.1 spike. Pseudovirus neutralization assays were performed as described previously (38) and in *Standard pseudovirus neutralization assays* section below. Before use, all sera were inactivated for 1 h at 56°C.

For each sample 1.5 million transcription units of spike pseudotyped library virus were mixed with RDPro pseudotyped virus at 1-2 % of total transcription units used. For each sera we performed selection at three concentrations aiming to neutralize more than 60% of library variants in at least two of these concentrations. Our starting serum dilution was twice the IC99 value as determined by standard pseudovirus neutralization, which typically significantly underestimates neutralization achieved for deep mutational scanning (perhaps due to differing amounts of spike on the surface of pseudoviruses used in standard neutralization assay versus library virus, or depletion of antibody molecules by the higher virion concentration in the library experiments). An example of neutralization achieved by different serum concentration can be seen here https://dms-vep.org/SARS-CoV-2_KP.3.1.1_spike_DMS/notebooks/avg_escape_antibody_escape_adult-1_pre_vaccination.html in the probability escape plots. Generally, serum escape probabilities > 0.4 allow identification of mutations that affect serum neutralization. Antibodies we used the following concentrations: BD55-1205 these concentrations were 0.73, 2.18, and 6.55 µg/ml, for SA55 0.32, 0.95, and 2.84 µg/ml, and for VYD222 100, 300, and 900 µg/ml. In standard pseudovirus neutralization assays all these concentrations were above IC99 value, but in deep mutational scanning data these ranged between IC50-IC99 for BD55-1205, IC5-IC75 for SA55 and IC94-IC99 for VYD222. After incubation, virus mixtures were used to infect medium-ACE2 cells in the presence of 2.5 µg/ml of amphotericin B and 15 h post infection non-integrated viral genomes were recovered and prepared for illumina sequencing as described previously (21).

To determine mutations which affect serum or antibody neutralization we used a biophysical model from *polyclonal* (v6.16) package (81), which is implemented in *dms-vep-pipeline-3* (v3.27.0) https://github.com/dms-vep/dms-vep-pipeline-3/tree/main.

Mean and individual sera escape plots and links to raw numeric escape values for each sera are at https://dms-vep.org/SARS-CoV-2_KP.3.1.1_spike_DMS/polyclonal_sera_escape.html. Interactive plots showing escape for BD55-1205, SA55 and VYD22 antibodies are at https://dms-vep.org/SARS-CoV-2_KP.3.1.1_spike_DMS/antibody_escape.html. The latter link also contains interactive structure visualizations showing deep mutational scanning measured escape in the context of RBD bound to each of the antibodies.

### Estimate of mutation effects on RBD up/down motion

To quantify a site’s effect on RBD up/down motion we used the following formula:

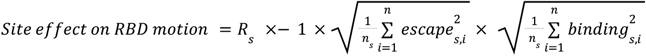

where ***R*** is Pearson correlation between mutation effects on serum escape (averaged across all sera) and ACE2 binding for site *s*. Positive R values were set to zero and then R was multiplied by negative 1. The root mean square of mutation effects on serum escape is calculated as 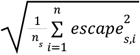, where *escape*_*s,I*_ is the measured serum escape effect (averaged across all sera) of mutation *n* at site *s*, and *n*_*s*_ is the number of mutations measured at site *s*, and 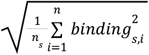 is the root mean square of mutation effects on ACE2 binding.

### Comparison with prior XBB.1.5 spike deep mutational scanning

Pseudovirus based deep mutational scanning data for XBB.1.5 spike was published previously in Dadonaite et al (2024). That dataset included two spike libraries: full spike deep mutational scanning library, where a subset of mutations was included throughout the spike protein, and RBD-only library, where all possible mutations were introduced only in the RBD. **Fig. 3C** compares ACE2 binding data for KP.3.1.1 deep mutational scanning libraries with full spike XBB.1.5 libraries and **Fig. 3D** compares ACE2 binding data for KP.3.1.1 deep mutational scanning libraries and XBB.1.5 RBD-only libraries for sites 435 and 493.

### Standard pseudovirus neutralization assays

Desired mutations were cloned into KP.3.1.1 spike expression plasmid https://github.com/dms-vep/SARS-CoV-2_KP.3.1.1_spike_DMS/blob/main/KP311_validation_notebooks/plasmid_maps/HDM_KP.3.1.1.gb and sequence was confirmed using whole plasmid sequencing. Spike pseudotyped lentiviruses were rescued by transfecting 293T cells with spike expression plasmids, Gag/Pol (BEI: NR-52517) helper plasmid and pHAGE6_Luciferase_IRES_ZsGreen backbone. 48 hours post transfection virus-containing cell supernatants were collected and titrated. Neutralization assays were performed as described in Crawford et al. (2020) using medium-ACE2 cells (24) in the presence of 2.5 µg/ml of amphotericin B. For all neutralization assays starting dilution was 0.05 and we performed eight 3-fold serial dilutions. Fraction infectivity at each dilution was determined relative to serum free controls and *neutcurve* (V2.1.0) package (82) was used to fit Hill curves to fraction infectivity data.

### Antibody Production

Antibodies were ordered from Genscript Biotech using published variable sequences (12,43,44,83). Variable sequences and complete expressed polypeptide sequences are specified in **Table S2**. These sequences were codon-optimized, cloned into expression vectors, and expressed in Chinese hamster ovary-derived cells. Heavy chain variable sequences were cloned into a human IgG1 backbone. The light chain variable sequences for BD55-1205 and SA55 were cloned into a human kappa light chain backbone; VYD222 was cloned into a human lambda light chain backbone.

### Cells

293T, 293T-rtTA, medium-ACE2 and cell-stored library cells were all grown in D10 media (Dulbecco’s Modified Eagle Medium with 10% heat-inactivated fetal bovine serum, 2 mM l-glutamine, 100 U/mL penicillin, and 100 μg/mL streptomycin). For deep mutational scanning library and 293T-rtTA cells tetracycline-free FBS was used. Medium-ACE2 cells were grown in the presence of 2 µg/ml doxycycline, which induced ACE2 expression in these cells.

### Ethics statement

Pre- and post-vaccination or infection sera were collected with informed consent from participants in the prospective longitudinal Hospitalized or Ambulatory Adults with Respiratory Viral Infections (HAARVI) study. The study was approved by University of Washington Institutional Review Board (#STUDY00000959).

## Supplementary figures

**Supplementary Figure 1.**
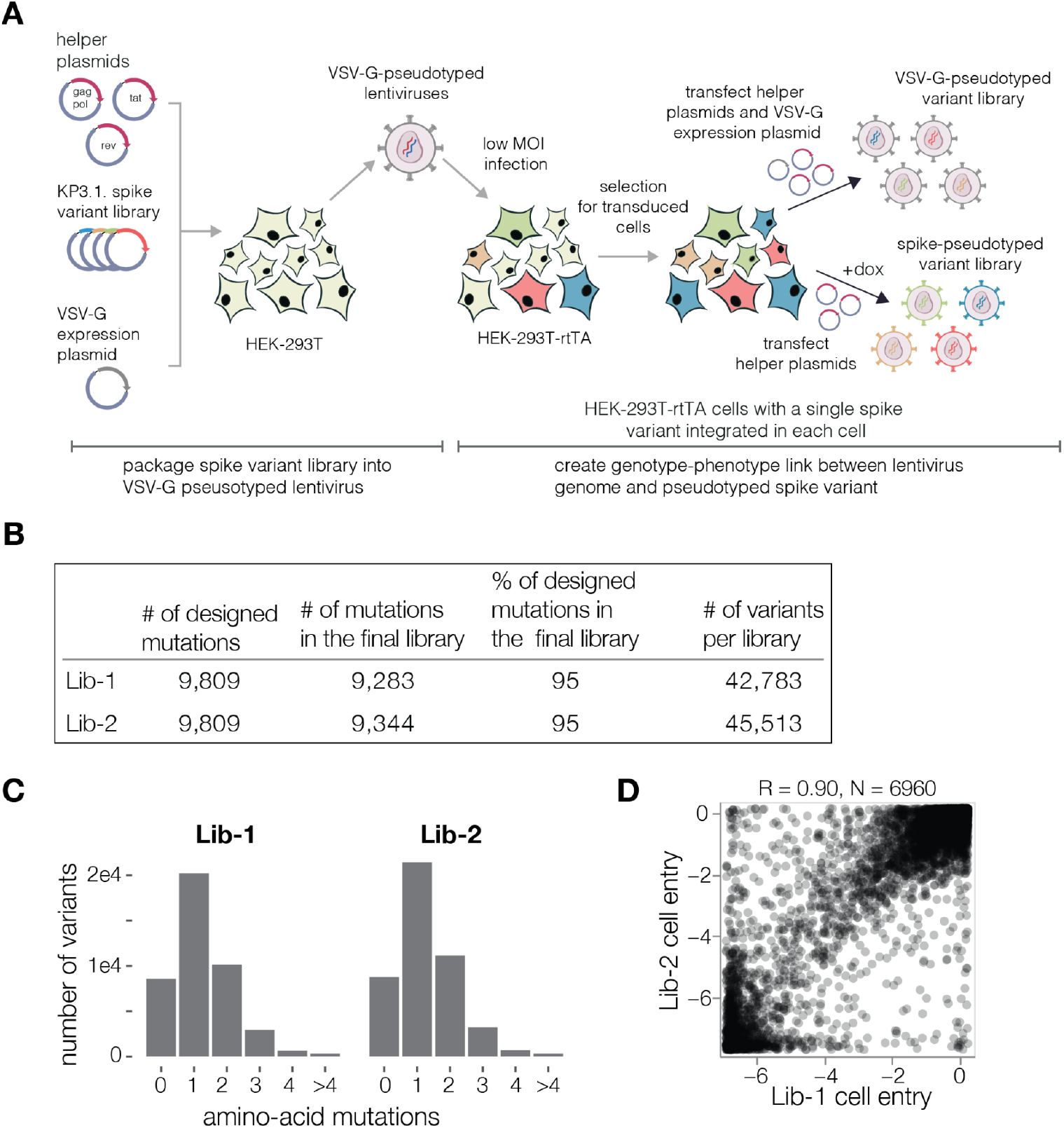
Design of KP.3.1.1 spike deep mutational scanning libraries. **A**. Method for producing genotype-phenotype linked pseudovirus-based deep mutational scanning libraries as applied to the KP.3.1.1 spike. 293T cells are transfected with lentivirus helper plasmids, lentiviral backbone plasmids encoding the barcoded KP.3.1.1 spike variant library, and VSV-G expression plasmid to produce VSV-G pseudotyped viruses. The viruses are then used to infect 293T cells expressing reverse tetracycline transactivator (rtTA) at low multiplicity of infection (MOI, <0.01) so that only a single virus genome integrates in any given cell. Cells are then selected for successful transduction using puromycin. From selected cells, genotype-phenotype linked virus libraries are made by inducing spike expression using doxycycline and transfecting lentivirus helper plasmids. To quantify the presence of non-functional as well as functional spike variants present in the libraries, we also rescue VSV-G pseudotyped viruses from the same library cells by transfecting lentivirus helper plasmids and VSV-G expression plasmids. **B**. Number of targeted and successfully included mutations in each of the two independent libraries. Note that our library design primarily targeted mutations expected to be functionally tolerated, see text and Methods for details. **C**. Distribution of mutations per spike variant for each library. **D**. Correlation between the cell entry effects for all high-confidence measured mutations in both of the two independent libraries. Throughout the paper we show the average measurement across both libraries unless otherwise indicated.

**Supplementary Figure 2.**
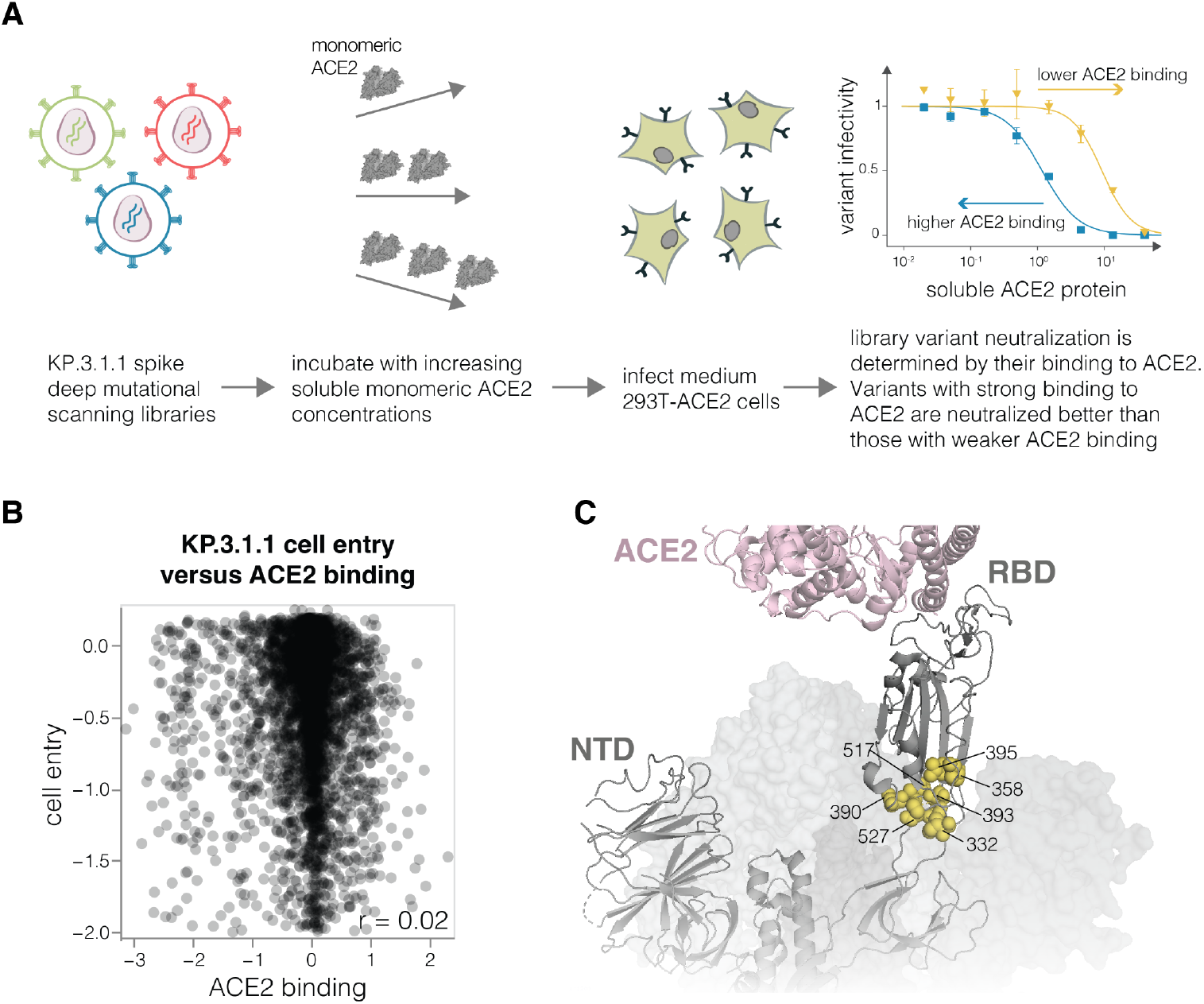
Effects of mutations on ACE2 binding. **A**. To measure effects of mutations to spike protein on ACE2 binding, deep mutational scanning libraries are incubated with monomeric soluble human ACE2 at multiple concentrations followed by infection of 293T-ACE2 cells expressing medium levels of ACE2. Library variants with mutations that increase ACE2 are better neutralized by soluble ACE2 compared to variants with mutations that decrease ACE2 binding. **B**. Correlation between mutation effects on ACE2 binding and cell entry; note it is only possible for our method to measure ACE2 binding for mutations that maintain at least modest levels of cell entry. **C**. SARS-CoV-2 spike structure with one RBD up in contact with ACE2 (PDB: 8IOU). Spheres show ACE2 distal RBD sites with strong effects on ACE2 binding as highlighted in **Fig. 3B**.

**Supplementary Figure 3.**
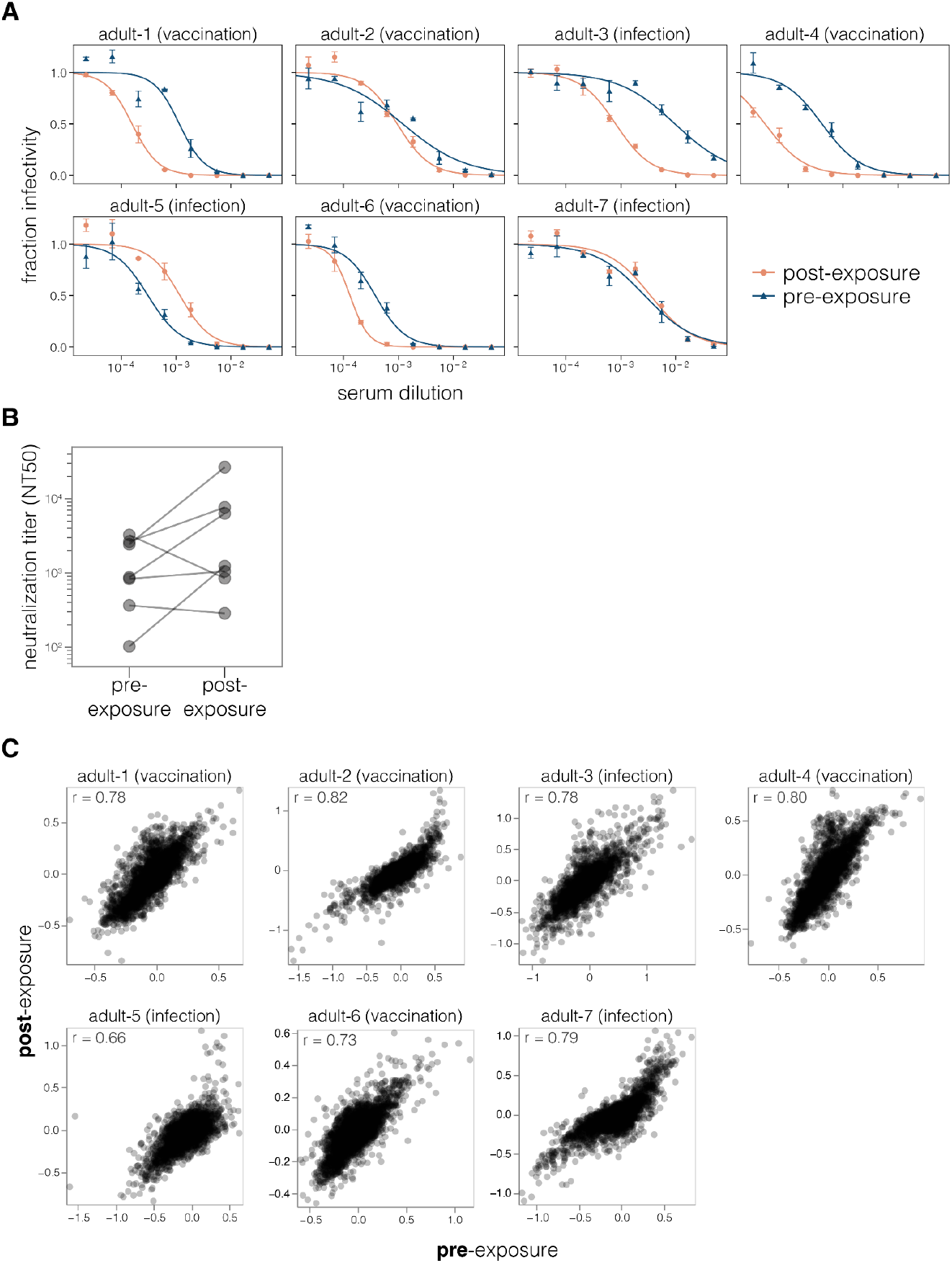
Neutralization of KP.3.1.1 pseudovirus by serum from the same individual collected pre- and post-exposure to a JN.1 descendant spike. **A**. Neutralization curves for the sera from seven individuals analyzed in this study. Each plot shows sera from the same individual pre- and post-exposure to JN.1-descendant spike. Curves were measured using standard neutralization assays with KP.3.1.1 spike pseudotyped lentiviral particles. **B**. Neutralizing titers against KP.3.1.1 pseudovirus for sera pre- and post-exposure to JN.1-descendant spike calculated from the neutralization curves in A. **C**. Correlation between deep mutational scanning measured sera escape scores for pre- and post-JN.1-descendant spike exposed sera for each individual. Each point is a different mutation and shows the measured effect of that mutation on escape from each serum.

**Supplementary Figure 4.**
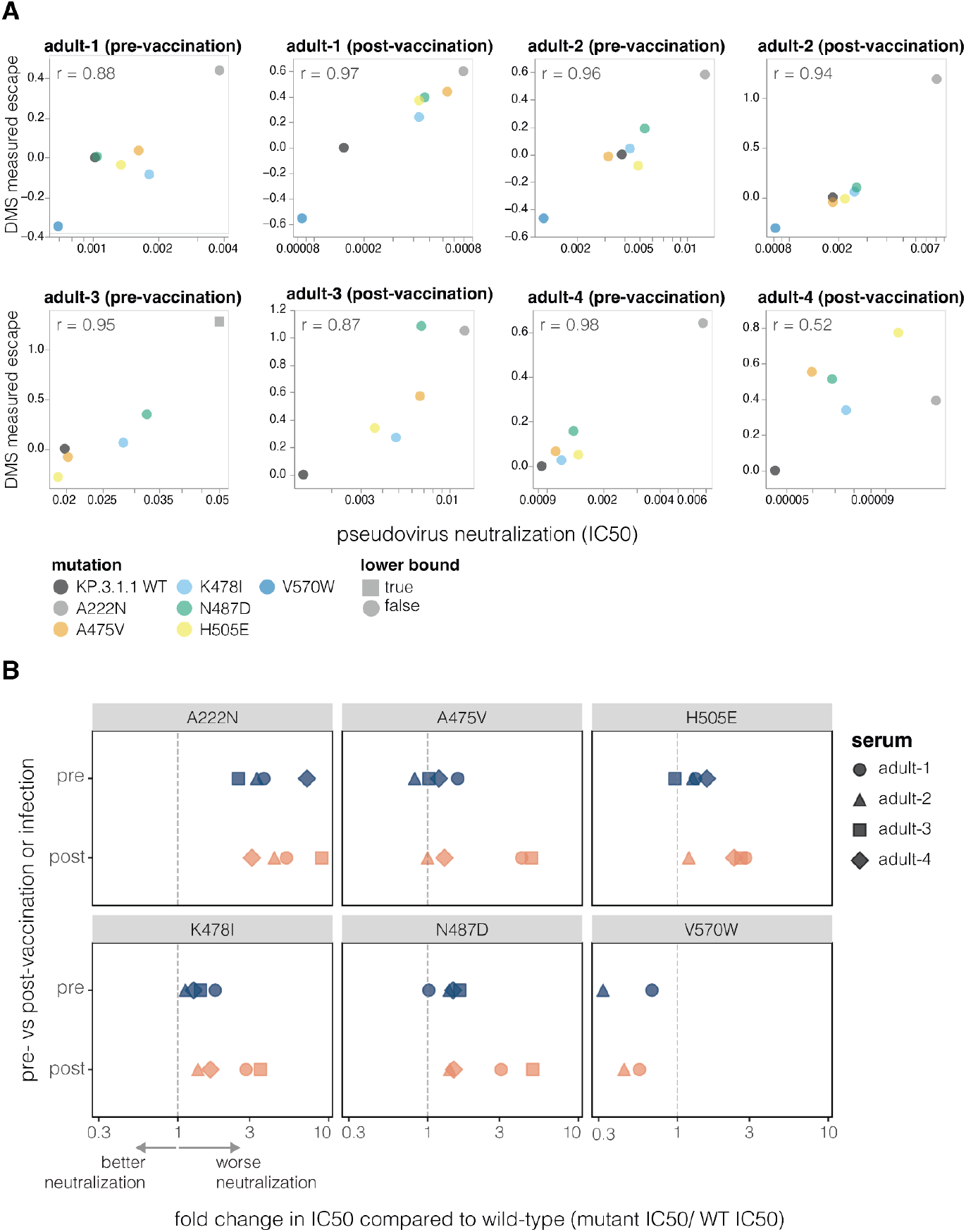
Effects of mutations to KP.3.1.1 spike on serum neutralization as measured by pseudovirus neutralization assay. **A**. Correlation between deep mutational scanning measured escape scores and IC50 values measured using a standard pseudovirus neutralization assay for various KP.3.1.1 spike mutants. **B**. Fold change in IC50 values for different KP.3.1.1 spike mutants relative to the unmutated KP.3.1.1 spike for pre- and post-vaccination or infection sera, as measured using a standard pseudovirus neutralization assay. All mutations were measured for four sera except for V570W which was only measured for two sera.

**Supplementary Figure 5.**
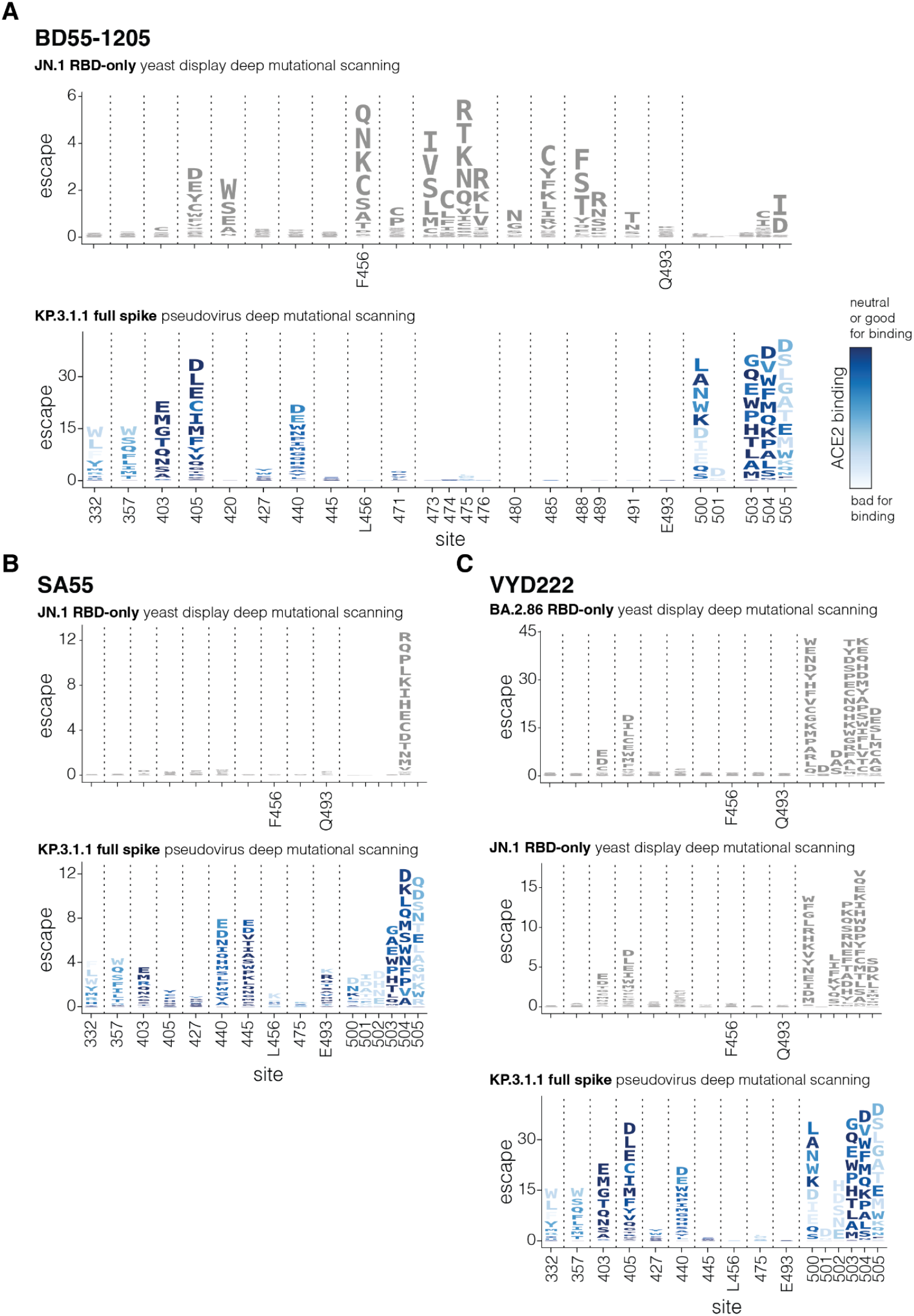
Escape for BD55-1205, SA55 and VYD222 antibodies measured by yeast-based RBD versus pseudovirus-based full-spike deep mutational scanning. **A**. Top logoplot shows effects of mutations at key sites on BD55-1205 antibody binding as measured using yeast-based deep mutational scanning of JN.1 RBD in previously published work by Jian et al (6). The height of the letter indicates binding escape for each mutation. The bottom logoplot shows mutations effects on neutralization by BD55-1205 as measured by KP.3.1.1 full-spile deep mutational scanning and is the same as in **Fig. 6**, with mutations colored according to their effect on ACE2 binding in the KP.3.1.1 full-spike pseudovirus deep mutational scanning. Sites where the parental amino acid differs between JN.1 and KP.3.1.1 backgrounds are labeled in both logoplots; sites labeled in just the bottom logoplot have the same parental amino acid in both JN.1 and KP.3.1.1. **B**. Same as A but for SA55 antibody. **C**. Same as A and B but for VYD222 antibody and with additional escape data for BA.2.86 RBD from Yuan et al (45) yeast-based RBD deep mutational scanning. No measurement was reported for site 502 for BD55-1205, and sites 502 and 505 for SA55 in the Jian et al RBD-only deep mutational scanning data. For BD55-1205 no measurements are available for sites 480 and 488 in full spike deep mutational scanning because all mutations at those sites are highly deleterious for cell entry.

**Supplementary Figure 6.**
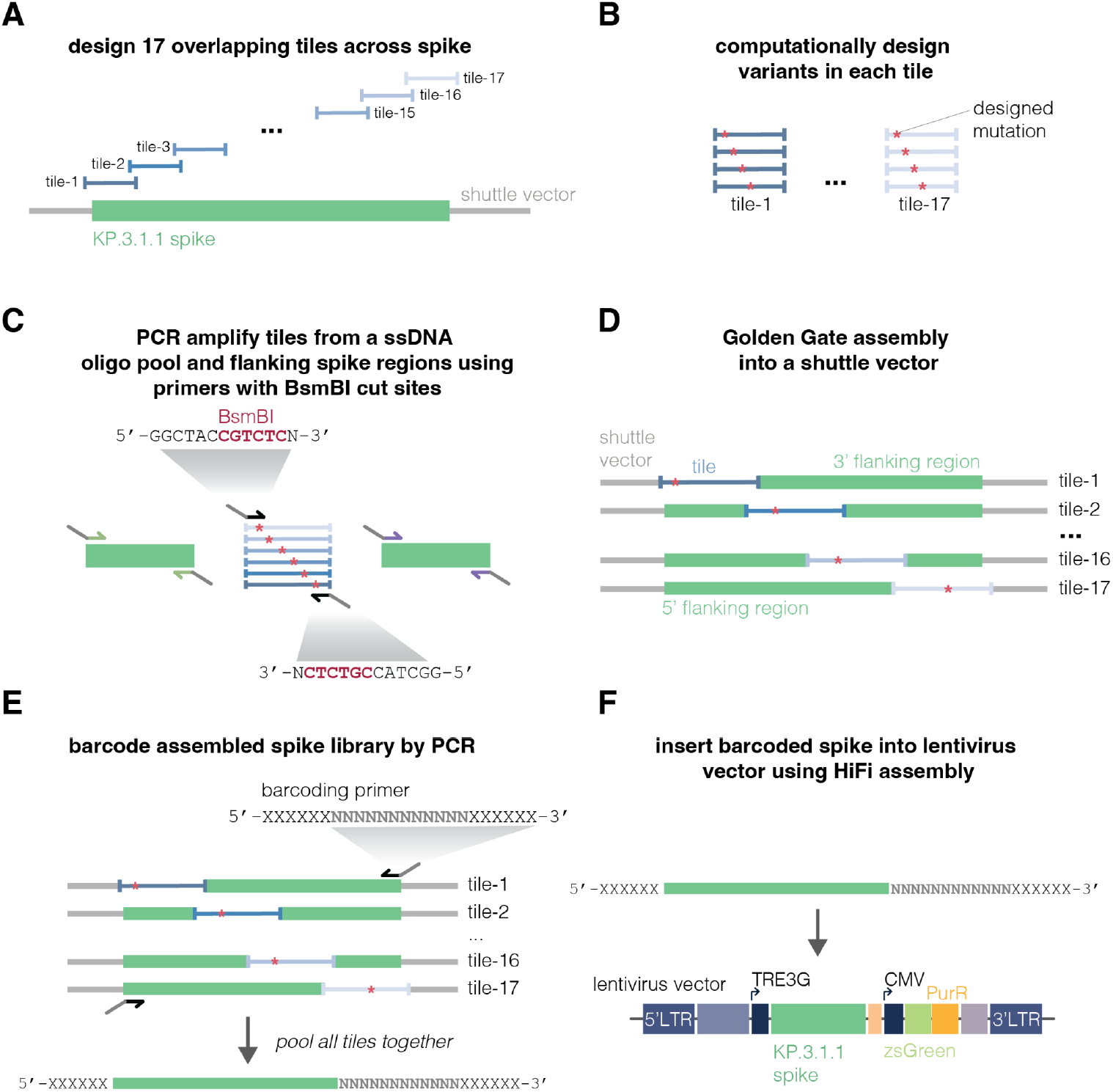
Cloning of KP.3.1.1 deep mutational scanning library. **A**. To produce plasmid library for deep mutational scanning using Golden Gate assembly, the KP.3.1.1 spike sequence was divided into 17 overlapping tiles. **B**. For each tile we computationally designed a pool of oligos containing all desired mutations. **C**. Designed oligos were ordered as a single-stranded DNA (ssDNA) oligo pool from which oligos belonging to each of the 17 tiles were amplified with primers containing the BsmBI restriction site. Unmutated spike sequences flanking each tile were also amplified. **D**. Golden Gate assembly was performed to assemble each tile pool and flanking spike sequences into a shuttle vector. **E**. Assembled spike sequences were amplified and barcoded in the same PCR reaction. Amplified and barcoded spike sequences were pooled equimolarly to make one library. **F**. The barcoded spike pool was cloned into a lentiviral vector using a HiFi reaction.

## Supplementary Tables

**Supplementary Table 1. Information about sera used in this study**

We used pre- and post-exposure sera from seven adults. The table indicates the last exposure (by vaccination or infection) and the days after this exposure that the “post-exposure” sera was collected. The “pre-exposure” serum from each individual was the last blood drawn prior to this final exposure, although individuals had different number of vaccinations or infections before this “pre-exposure” serum was collected. The table also indicates the last exposure before the “pre-exposure” serum collection. In all cases, the “post-exposure” serum was after a vaccination with the KP.2 spike or an infection in May-November of 2024, when JN.1-descendant variants dominated in Washington state where all the sera were collected (84).

**Supplementary Table 2. Antibody sequences**

Variable chain sequences for BD55-1205, SA55 and VYD222 antibodies. The complete expressed polypeptide sequences, including the human IgG1, lambda, or kappa constant sequences are provided. Heavy (HC) and light chains (LC) were expressed with murine Ig heavy chain V region 102 as an N-terminal export signal sequence.

## References

1. Bloom JD, Neher RA. Fitness effects of mutations to SARS-CoV-2 proteins. Virus Evol. 2023 Jan 1;9(2):vead055.

2. Dadonaite B, Brown J, McMahon TE, Farrell AG, Figgins MD, Asarnow D, et al. Spike deep mutational scanning helps predict success of SARS-CoV-2 clades. Nature. 2024 July;631(8021):617–26.

3. Lamb KD, Luka MM, Saathoff M, Orton RJ, Phan MVT, Cotten M, et al. Mutational signature dynamics indicate SARS-CoV-2’s evolutionary capacity is driven by host antiviral molecules. PLOS Comput Biol. 2024 Jan 25;20(1):e1011795.

4. Greaney AJ, Starr TN, Bloom JD. An antibody-escape estimator for mutations to the SARS-CoV-2 receptor-binding domain. Virus Evol. 2022 Jan 1;8(1):veac021.

5. Greaney AJ, Loes AN, Gentles LE, Crawford KHD, Starr TN, Malone KD, et al. Antibodies elicited by mRNA-1273 vaccination bind more broadly to the receptor binding domain than do those from SARS-CoV-2 infection. Sci Transl Med. 2021 June 30;13(600):eabi9915.

6. Jian F, Wang J, Yisimayi A, Song W, Xu Y, Chen X, et al. Evolving antibody response to SARS-CoV-2 antigenic shift from XBB to JN.1. Nature. 2024 Nov 7;1–3.

7. Feng L, Sun Z, Zhang Y, Jian F, Yang S, Xia K, et al. Structural and molecular basis of the epistasis effect in enhanced affinity between SARS-CoV-2 KP.3 and ACE2. Cell Discov. 2024 Nov 30;10(1):123.

8. Starr TN, Greaney AJ, Hannon WW, Loes AN, Hauser K, Dillen JR, et al. Shifting mutational constraints in the SARS-CoV-2 receptor-binding domain during viral evolution. Science. 2022 July 22;377(6604):420–4.

9. Starr TN, Greaney AJ, Hilton SK, Ellis D, Crawford KHD, Dingens AS, et al. Deep Mutational Scanning of SARS-CoV-2 Receptor Binding Domain Reveals Constraints on Folding and ACE2 Binding. Cell. 2020 Sept 3;182(5):1295-1310.e20.

10. Ozono S, Zhang Y, Ode H, Sano K, Tan TS, Imai K, et al. SARS-CoV-2 D614G spike mutation increases entry efficiency with enhanced ACE2-binding affinity. Nat Commun. 2021 Feb 8;12(1):848.

11. Khatri R, Siddqui G, Sadhu S, Maithil V, Vishwakarma P, Lohiya B, et al. Intrinsic D614G and P681R/H mutations in SARS-CoV-2 VoCs Alpha, Delta, Omicron and viruses with D614G plus key signature mutations in spike protein alters fusogenicity and infectivity. Med Microbiol Immunol (Berl). 2023 Feb 1;212(1):103–22.

12. Jian F, Wec AZ, Feng L, Yu Y, Wang L, Wang P, et al. Viral evolution prediction identifies broadly neutralizing antibodies to existing and prospective SARS-CoV-2 variants. Nat Microbiol. 2025 June 10;1–15.

13. Lei R, Qing E, Odle A, Yuan M, Gunawardene CD, Tan TJC, et al. Functional and antigenic characterization of SARS-CoV-2 spike fusion peptide by deep mutational scanning. Nat Commun. 2024 May 14;15:4056.

14. Tan TJC, Mou Z, Lei R, Ouyang WO, Yuan M, Song G, et al. High-throughput identification of prefusion-stabilizing mutations in SARS-CoV-2 spike. Nat Commun. 2023 Apr 10;14(1):2003.

15. Taylor AL, Starr TN. Deep mutational scanning of SARS-CoV-2 Omicron BA.2.86 and epistatic emergence of the KP.3 variant. Virus Evol. 2024 Oct 17;10(1):veae067.

16. Jian F, Feng L, Yang S, Yu Y, Wang L, Song W, et al. Convergent evolution of SARS-CoV-2 XBB lineages on receptor-binding domain 455–456 synergistically enhances antibody evasion and ACE2 binding. PLOS Pathog. 2023 Dec 20;19(12):e1011868.

17. Meijers M, Ruchnewitz D, Eberhardt J, Łuksza M, Lässig M. Population immunity predicts evolutionary trajectories of SARS-CoV-2. Cell. 2023 Nov 9;186(23):5151-5164.e13.

18. Suntronwong N, Assawakosri S, Klinfueng S, Duangchinda T, Chantima W, Pakchotanon P, et al. Age associated SARS-CoV-2 immune responses provide insights into population immunity over four years since the COVID-19 pandemic. Sci Rep. 2025 July 2;15(1):23183.

19. Raharinirina NA, Gubela N, Börnigen D, Smith MR, Oh DY, Budt M, et al. SARS-CoV-2 evolution on a dynamic immune landscape. Nature. 2025 Mar;639(8053):196–204.

20. Dadonaite B, Burrell AR, Logue J, Chu HY, Payne DC, Haslam DB, et al. SARS-CoV-2 neutralizing antibody specificities differ dramatically between recently infected infants and immune-imprinted individuals. J Virol. 2025 Mar 25;99(4):e00109–25.

21. Dadonaite B, Crawford KHD, Radford CE, Farrell AG, Yu TC, Hannon WW, et al. A pseudovirus system enables deep mutational scanning of the full SARS-CoV-2 spike. Cell. 2023 Mar 16;186(6):1263-1278.e20.

22. Roemer, Cornelius, Neher, Richard. GitHub. 2024 [cited 2025 July 2]. SARS-CoV-2_variant-reports_2024-12-20. Available from: https://github.com/neherlab/SARS-CoV-2_variant-reports/blob/b85f7caeb1503ad2a803d3e38568a4e3c767216c/reports/variant_report_2024-12-20.md

23. World Health Organisation. Statement on the antigen composition of COVID-19 vaccines [Internet]. 2025 [cited 2025 July 2]. Available from: https://www.who.int/news/item/15-05-2025-statement-on-the-antigen-composition-of-covid-19-vaccines

24. Farrell AG, Dadonaite B, Greaney AJ, Eguia R, Loes AN, Franko NM, et al. Receptor-Binding Domain (RBD) Antibodies Contribute More to SARS-CoV-2 Neutralization When Target Cells Express High Levels of ACE2. Viruses. 2022 Sept;14(9):2061.

25. Cantoni D, Murray MJ, Kalemera MD, Dicken SJ, Stejskal L, Brown G, et al. Evolutionary remodelling of N-terminal domain loops fine-tunes SARS-CoV-2 spike. EMBO Rep. 2022 Oct 6;23(10):e54322.

26. Bayati A, Kumar R, Francis V, McPherson PS. SARS-CoV-2 infects cells after viral entry via clathrin-mediated endocytosis. J Biol Chem. 2021 Jan 1;296:100306.

27. Jackson CB, Farzan M, Chen B, Choe H. Mechanisms of SARS-CoV-2 entry into cells. Nat Rev Mol Cell Biol. 2022 Jan;23(1):3–20.

28. Cao Y, Jian F, Wang J, Yu Y, Song W, Yisimayi A, et al. Imprinted SARS-CoV-2 humoral immunity induces convergent Omicron RBD evolution. Nature. 2023 Feb;614(7948):521–9.

29. Wang Q, Iketani S, Li Z, Guo Y, Yeh AY, Liu M, et al. Antigenic characterization of the SARS-CoV-2 Omicron subvariant BA.2.75. Cell Host Microbe. 2022 Nov 9;30(11):1512-1517.e4.

30. Wang Q, Nair MS, Anang S, Zhang S, Nguyen H, Huang Y, et al. Functional differences among the spike glycoproteins of multiple emerging severe acute respiratory syndrome coronavirus 2 variants of concern. iScience. 2021 Nov 19;24(11):103393.

31. Walls AC, Xiong X, Park YJ, Tortorici MA, Snijder J, Quispe J, et al. Unexpected Receptor Functional Mimicry Elucidates Activation of Coronavirus Fusion. Cell. 2019 Feb 21;176(5):1026-1039.e15.

32. Tse AL, Acreman CM, Ricardo-Lax I, Berrigan J, Lasso G, Balogun T, et al. Distinct pathways for evolution of enhanced receptor binding and cell entry in SARS-like bat coronaviruses. PLOS Pathog. 2024 Nov 15;20(11):e1012704.

33. Uğurlu H, Salminen P, Yuan M, Fagerlund R, Mäkelä AR, Žusinaite E, et al. Altered spike dynamics drives immune evasion of SARS-CoV-2 [Internet]. bioRxiv; 2024 [cited 2025 July 19]. p. 2024.12.03.626570. Available from: https://www.biorxiv.org/content/10.1101/2024.12.03.626570v1

34. Taylor AL, Starr TN. Deep mutational scans of XBB.1.5 and BQ.1.1 reveal ongoing epistatic drift during SARS-CoV-2 evolution. PLOS Pathog. 2023 Dec 29;19(12):e1011901.

35. Wang Q, Mellis IA, Ho J, Bowen A, Kowalski-Dobson T, Valdez R, et al. Recurrent SARS-CoV-2 spike mutations confer growth advantages to select JN.1 sublineages. Emerg Microbes Infect. 13(1):2402880.

36. Yajima H, Anraku Y, Kaku Y, Kimura KT, Plianchaisuk A, Okumura K, et al. Structural basis for receptor-binding domain mobility of the spike in SARS-CoV-2 BA.2.86 and JN.1. Nat Commun. 2024 Oct 7;15(1):8574.

37. Wang Q, Guo Y, Mellis IA, Wu M, Mohri H, Gherasim C, et al. Antibody evasiveness of SARS-CoV-2 subvariants KP.3.1.1 and XEC. Cell Rep. 2025 Apr 22;44(4):115543.

38. Crawford KHD, Eguia R, Dingens AS, Loes AN, Malone KD, Wolf CR, et al. Protocol and Reagents for Pseudotyping Lentiviral Particles with SARS-CoV-2 Spike Protein for Neutralization Assays. Viruses. 2020 May 6;12(5):513.

39. Hodcroft EB, Zuber M, Nadeau S, Vaughan TG, Crawford KHD, Althaus CL, et al. Spread of a SARS-CoV-2 variant through Europe in the summer of 2020. Nature. 2021 July;595(7869):707–12.

40. Zhao Z, Zhou J, Tian M, Huang M, Liu S, Xie Y, et al. Omicron SARS-CoV-2 mutations stabilize spike up-RBD conformation and lead to a non-RBM-binding monoclonal antibody escape. Nat Commun. 2022 Aug 24;13(1):4958.

41. Yang H, Guo H, Wang A, Cao L, Fan Q, Jiang J, et al. Structural basis for the evolution and antibody evasion of SARS-CoV-2 BA.2.86 and JN.1 subvariants. Nat Commun. 2024 Sept 4;15(1):7715.

42. Zhang QE, Lindenberger J, Parsons RJ, Thakur B, Parks R, Park CS, et al. SARS-CoV-2 Omicron XBB lineage spike structures, conformations, antigenicity, and receptor recognition. Mol Cell. 2024 July 25;84(14):2747-2764.e7.

43. Cao Y, Jian F, Zhang Z, Yisimayi A, Hao X, Bao L, et al. Rational identification of potent and broad sarbecovirus-neutralizing antibody cocktails from SARS convalescents. Cell Rep. 2022 Dec 20;41(12):111845.

44. Ison MG, Popejoy M, Evgeniev N, Tzekova M, Mahoney K, Betancourt N, et al. Efficacy and Safety of Adintrevimab (ADG20) for the Treatment of High-Risk Ambulatory Patients With Mild or Moderate Coronavirus Disease 2019: Results From a Phase 2/3, Randomized, Placebo-Controlled Trial (STAMP) Conducted During Delta Predominance and Early Emergence of Omicron. Open Forum Infect Dis. 2023 May 24;10(6):ofad279.

45. Yuan M, West BR, Foreman WB, Powers C, Feng Z, Taylor AL, et al. Structural and functional analysis of VYD222: a broadly neutralizing antibody against SARS-CoV-2 variants [Internet]. bioRxiv; 2025 [cited 2025 Sept 12]. p. 2025.08.28.672883. Available from: https://www.biorxiv.org/content/10.1101/2025.08.28.672883v1

46. SINOVAC Reports on the Clinical Trial Progress of its Anti-COVID-19 Antibody SA55 Injection-SINOVAC: Supply Vaccines to Eliminate Human Diseases [Internet]. [cited 2025 July 25]. Available from: https://www.sinovac.com/en-us/news/SINOVAC_Reports_on_the_Clinical_Trial_Progress_of_its_Anti_COVID_19_Antibody_SA55_Injection

47. Invivyd. Invivyd Announces FDA Authorization for Emergency Use of PEMGARDATM (Formerly VYD222) for Pre-exposure Prophylaxis (PrEP) of COVID-19 - Invivyd [Internet]. [cited 2025 July 23]. Available from: https://investors.invivyd.com/news-releases/news-release-details/invivyd-announces-fda-authorization-emergency-use-pemgardatm/

48. Cao Y, Wang J, Jian F, Xiao T, Song W, Yisimayi A, et al. Omicron escapes the majority of existing SARS-CoV-2 neutralizing antibodies. Nature. 2022 Feb;602(7898):657–63.

49. Rappazzo CG, Tse LV, Kaku CI, Wrapp D, Sakharkar M, Huang D, et al. Broad and potent activity against SARS-like viruses by an engineered human monoclonal antibody. Science. 2021 Feb 19;371(6531):823–9.

50. Powers C, Williams B, Kreher A, Gao F, West B, Chupp D, et al. Neutralization of recent SARS-CoV-2 variants by genetically and structurally related mAbs of the pemivibart lineage [Internet]. bioRxiv; 2024 [cited 2025 July 23]. p. 2024.11.11.623127. Available from: https://www.biorxiv.org/content/10.1101/2024.11.11.623127v1

51. Wang Q, Guo Y, Ho J, Ho DD. Activity of Research-Grade Pemivibart against Recent SARS-CoV-2 JN.1 Sublineages. N Engl J Med. 2024 Nov 13;391(19):1863–4.

52. Liu L, Iketani S, Guo Y, Chan JFW, Wang M, Liu L, et al. Striking antibody evasion manifested by the Omicron variant of SARS-CoV-2. Nature. 2022 Feb;602(7898):676–81.

53. Tse AL, Lasso G, McLellan JS, Chandran K, Miller EH. Bat sarbecovirus WIV1-CoV bears an adaptive mutation that alters spike dynamics and enhances ACE2 binding [Internet]. bioRxiv; 2025 [cited 2025 Aug 8]. p. 2025.04.14.648681. Available from: https://www.biorxiv.org/content/10.1101/2025.04.14.648681v1

54. Li Q, Cai X, Li X, Zhang Y, Li R, Kang Z, et al. Structural and functional constraints on spike activation and host protease utilization limit cell entry of SARS-CoV-2-related bat coronaviruses. J Virol. 2025 July 24;0(0):e01007–25.

55. Greaney AJ, Loes AN, Crawford KHD, Starr TN, Malone KD, Chu HY, et al. Comprehensive mapping of mutations in the SARS-CoV-2 receptor-binding domain that affect recognition by polyclonal human plasma antibodies. Cell Host Microbe. 2021 Mar 10;29(3):463-476.e6.

56. Piccoli L, Park YJ, Tortorici MA, Czudnochowski N, Walls AC, Beltramello M, et al. Mapping Neutralizing and Immunodominant Sites on the SARS-CoV-2 Spike Receptor-Binding Domain by Structure-Guided High-Resolution Serology. Cell. 2020 Nov 12;183(4):1024-1042.e21.

57. Premkumar L, Segovia-Chumbez B, Jadi R, Martinez DR, Raut R, Markmann AJ, et al. The receptor-binding domain of the viral spike protein is an immunodominant and highly specific target of antibodies in SARS-CoV-2 patients. Sci Immunol. 2020 June 11;5(48):eabc8413.

58. Liang CY, Raju S, Liu Z, Li Y, Arunkumar GA, Case JB, et al. Imprinting of serum neutralizing antibodies by Wuhan-1 mRNA vaccines. Nature. 2024 June;630(8018):950–60.

59. Kaku CI, Starr TN, Zhou P, Dugan HL, Khalifé P, Song G, et al. Evolution of antibody immunity following Omicron BA.1 breakthrough infection. Nat Commun. 2023 May 12;14(1):2751.

60. Addetia A, Piccoli L, Case JB, Park YJ, Beltramello M, Guarino B, et al. Neutralization, effector function and immune imprinting of Omicron variants. Nature. 2023 Sept;621(7979):592–601.

61. Yisimayi A, Song W, Wang J, Jian F, Yu Y, Chen X, et al. Repeated Omicron exposures override ancestral SARS-CoV-2 immune imprinting. Nature. 2024 Jan;625(7993):148–56.

62. Schiepers A, van ‘t Wout MFL, Greaney AJ, Zang T, Muramatsu H, Lin PJC, et al. Molecular fate-mapping of serum antibody responses to repeat immunization. Nature. 2023 Mar;615(7952):482–9.

63. Roemer C, Neher RA. GitHub. 2025 [cited 2025 Aug 7]. SARS-CoV-2 variant reports. Available from: https://github.com/neherlab/SARS-CoV-2_variant-reports/blob/main/reports/variant_report_202 5-04-23.md

64. Barrat-Charlaix P, Neher RA. Eco-evolutionary dynamics of adapting pathogens and host immunity. eLife [Internet]. 2024 Oct 18 [cited 2025 Aug 8];13. Available from: https://elifesciences.org/reviewed-preprints/97350

65. Planas D, Staropoli I, Planchais C, Yab E, Jeyarajah B, Rahou Y, et al. Escape of SARS-CoV-2 Variants KP.1.1, LB.1, and KP3.3 From Approved Monoclonal Antibodies. Pathog Immun. 2024 Sept 30;10(1):1–11.

66. Guo C, Yu Y, Liu J, Jian F, Yang S, Song W, et al. Antigenic and virological characteristics of SARS-CoV-2 variants BA.3.2, XFG, and NB.1.8.1. Lancet Infect Dis. 2025 July 1;25(7):e374–7.

67. Khare S, Gurry C, Freitas L, Schultz MB, Bach G, Diallo A, et al. GISAID’s role in pandemic response. China CDC Wkly. 2021;3(49):1049.

68. Turakhia Y, Thornlow B, Hinrichs AS, De Maio N, Gozashti L, Lanfear R, et al. Ultrafast Sample placement on Existing tRees (UShER) enables real-time phylogenetics for the SARS-CoV-2 pandemic. Nat Genet. 2021 June;53(6):809–16.

69. O’Toole Á, Pybus OG, Abram ME, Kelly EJ, Rambaut A. Pango lineage designation and assignment using SARS-CoV-2 spike gene nucleotide sequences. BMC Genomics. 2022 Feb 11;23(1):121.

70. Hanning KR, Walker EJ, Beijerling K, Irvine EB, Steel JJ, Kelton W. Simple high-throughput encoding of deep mutational scanning libraries by oligo-based Golden Gate assembly [Internet]. bioRxiv; 2025 [cited 2025 Aug 12]. p. 2025.07.16.665225. Available from: https://www.biorxiv.org/content/10.1101/2025.07.16.665225v1

71. Jann J, Gagnon-Arsenault I, Pageau A, Dubé AK, Fijarczyk A, Durand R, et al. Making deep mutational scanning accessible: a cost-efficient approach to construct barcoded libraries for genes of any length [Internet]. bioRxiv; 2025 [cited 2025 Aug 12]. p. 2025.05.29.656836. Available from: https://www.biorxiv.org/content/10.1101/2025.05.29.656836v1

72. Álvarez-Rodríguez B, Velandia-Álvarez S, Toft C, Geller R. Mapping the mutational landscape of a full viral proteome reveals distinct profiles of mutation tolerability [Internet]. bioRxiv; 2024 [cited 2025 Aug 12]. p. 2024.03.07.583990. Available from: https://www.biorxiv.org/content/10.1101/2024.03.07.583990v2

73. Dolan P, Bakhache W, Orr W, McCormick L. Uncovering Structural Plasticity of Enterovirus A through Deep Insertional and Deletional Scanning [Internet]. Research Square; 2024 [cited 2025 Aug 12]. Available from: https://www.researchsquare.com/article/rs-3835307/v1

74. Macdonald CB, Nedrud D, Grimes PR, Trinidad D, Fraser JS, Coyote-Maestas W. DIMPLE: deep insertion, deletion, and missense mutation libraries for exploring protein variation in evolution, disease, and biology. Genome Biol. 2023 Feb 24;24(1):36.

75. Coyote-Maestas W, Nedrud D, Okorafor S, He Y, Schmidt D. Targeted insertional mutagenesis libraries for deep domain insertion profiling. Nucleic Acids Res. 2020 Jan 24;48(2):e11.

76. Nedrud D, Coyote-Maestas W, Schmidt D. A large-scale survey of pairwise epistasis reveals a mechanism for evolutionary expansion and specialization of PDZ domains. Proteins Struct Funct Bioinforma. 2021;89(8):899–914.

77. Potapov V, Ong JL, Kucera RB, Langhorst BW, Bilotti K, Pryor JM, et al. Comprehensive Profiling of Four Base Overhang Ligation Fidelity by T4 DNA Ligase and Application to DNA Assembly. ACS Synth Biol. 2018 Nov 16;7(11):2665–74.

78. Mateyko N, de Boer CG. Culture Wars: Empirically Determining the Best Approach for Plasmid Library Amplification. ACS Synth Biol. 2024 Aug 16;13(8):2328–34.

79. Haddox HK, Galloway JG, Dadonaite B, Bloom JD, Matsen FA, DeWitt WS. Jointly modeling deep mutational scans identifies shifted mutational effects among SARS-CoV-2 spike homologs [Internet]. bioRxiv; 2023 [cited 2023 Oct 4]. p. 2023.07.31.551037. Available from: https://www.biorxiv.org/content/10.1101/2023.07.31.551037v1

80. Otwinowski J, McCandlish DM, Plotkin JB. Inferring the shape of global epistasis. Proc Natl Acad Sci. 2018 Aug 7;115(32):E7550–8.

81. Yu TC, Thornton ZT, Hannon WW, DeWitt WS, Radford CE, Matsen FA IV, et al. A biophysical model of viral escape from polyclonal antibodies. Virus Evol. 2022 July 1;8(2):veac110.

82. Loes AN, Tarabi RAL, Huddleston J, Touyon L, Wong SS, Cheng SMS, et al. High-throughput sequencing-based neutralization assay reveals how repeated vaccinations impact titers to recent human H1N1 influenza strains. J Virol. 2024 Sept 24;98(10):e00689–24.

83. CDER Scientific Review. Emergency Use Authorization (EUA) for PEMGARDA Center for Drug Evaluation and Research (CDER) Review Memorandum [Internet]. U.S. Food & Drug Administration; 2024. Available from: https://www.fda.gov/media/181308/download?attachment

84. UW Virology COVID-19 Dashboard [Internet]. [cited 2023 Nov 7]. Available from: https://depts.washington.edu/labmed/covid19/#sequencing-information

